# Decompose and Recompose: Reference-Centered Multimodal Adaptation of Protein Language Models for Enzyme Turnover Prediction

**DOI:** 10.64898/2026.11.15.766718

**Authors:** Aocheng Ding, Tengfei Ma, Xuan Lin, Shan Lin, Xiangrong Liu, Leyi Wei

**Affiliations:** School of Informatics, Xiamen University, Xiamen, China; State Key Laboratory of Chemo and Biosensing, Hunan University, Changsha, China; College of Computer Science and Electronic Engineering, Hunan University, Changsha, China; College of Computer Science, Xiangtan University, Xiangtan, China; Faculty of Applied Sciences, Macao Polytechnic University, Macao SAR, China

## Abstract

Predicting enzyme turnover numbers (k_cat_) from protein sequences and reaction chemistry supports virtual screening, which requires distinguishing activity differences between enzyme–reaction pairs. Recent interaction models face two representation challenges. First, they merge reactants and products before enzyme interaction, leaving separate enzyme responses implicit despite the two sides’ distinct roles in recognition, transformation, and release. Second, cross-attention readouts mix reaction-dependent changes with an enzyme-specific residue background, which can obscure activity-relevant differences across samples. To address these challenges, we introduce EnzRef, a lightweight framework for multimodal adaptation of frozen protein language models. EnzRef decomposes reaction chemistry into reactant and product conditions, models their interactions with the enzyme separately, and recomposes the resulting responses for activity prediction. During these interactions, Reference-Centered Cross-Attention expresses each response as a deviation from the enzyme’s own residue background to expose chemically conditioned differences for activity discrimination. With 1.72M trainable parameters, EnzRef achieves state-of-the-art source prediction on CatPred-DB and zero-shot transfer to TurNuP-DB. Across ten independent runs, EnzRef achieves 19.8% higher mean R^2^ than the strongest baseline MCKcat in zero-shot transfer to unseen enzymes. It also predicts activity differences more accurately between pairs sharing an enzyme or a reaction.

## 1. Introduction

Enzyme turnover prediction estimates *k*_cat_, the catalytic rate under substrate saturation, from an enzyme sequence and reaction chemistry. Computational screening can prioritize enzymes for experimental testing [1, 2], and activity-guided engineering depends on distinguishing the relative performance of candidates [3–5]. The same enzyme can exhibit different activities across reactions, while different enzymes can catalyze the same reaction at different rates. Useful multimodal representations must preserve these differences even when samples share an enzyme or a reaction.

Protein language models (PLMs) provide transferable enzyme representations [6–8], enriched by structure-aware and enzyme-focused pretraining [9–11]. Task adaptation connects these features to functional prediction through fine-tuning or task-specific modules [12–15]. Yet sequence information alone cannot distinguish reactions catalyzed by the same enzyme. Turnover predictors therefore combine protein features with substrate information [16, 17], complete reactant sets [18], or whole reactions [19]. When protein summaries are computed independently of chemistry, the same representation must serve different chemical conditions. Chemically conditioned interactions let molecular context shape the enzyme features used for prediction [20]. Recent methods such as PMAK [21] and MCKcat [22] use joint RXNFP embeddings [23] to condition residue aggregation.

Two representation challenges remain. First, reactants and products play distinct roles in catalysis: reactants are recognized and transformed, whereas products are formed and released [24, 25]. Recent residue-level interaction models such as PMAK and MCKcat nevertheless merge reactants and products into a single RXNFP embedding [23] before enzyme interaction, leaving the enzyme’s separate role responses and the relation between them implicit (Fig. 1a). Second, cross-attention readouts mix chemically conditioned changes with an enzyme-specific residue background. This background is shared across chemical conditions within an enzyme but varies across enzymes. Comparing raw readouts across samples therefore mixes variation in enzyme background with variation in chemical response, which can obscure activity-relevant distinctions (Fig. 1b).

**Figure 1.**
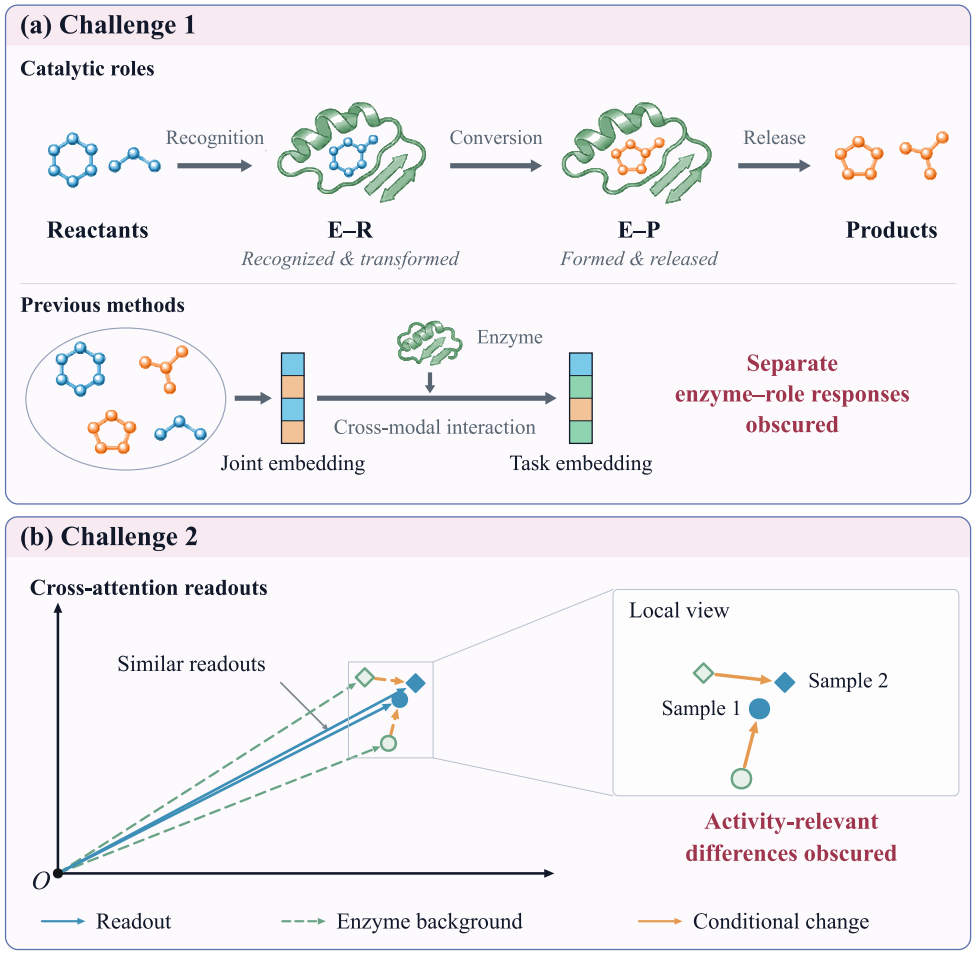
Two representation challenges in enzyme–reaction modeling. (a) Reactants and products play distinct catalytic roles, but joint reaction conditioning leaves their separate enzyme responses implicit. (b) Enzyme background can obscure conditional differences across samples with similar cross-attention readouts. The inset magnifies these conditional changes.

Reactant and product chemistry provide separate conditions for reading the same enzyme. Both readouts lie in the linear manifold spanned by its learned residue values, whose uniform average provides a shared, query-independent reference. Each attention-weighted average can then be expressed as this reference plus a query-dependent deviation. These deviations and their linearly recomposed components remain within this manifold. This linear recomposition preserves the information in both referenced responses while organizing their relation for learning activity differences across enzyme–reaction pairs.

Building on this view, we introduce ENZREF (Enzyme-Referenced Role Representations), a lightweight framework for multimodal adaptation of frozen PLM features. Chemical Semantic Decomposition (CSD) supplies separate reactant and product conditions for enzyme interaction, and Symmetry-Aware Semantic Recomposition (SASR) recomposes the resulting responses through their mean deviation and signed role contrast for activity prediction. Between these stages, Reference-Centered Cross-Attention (RefCA) expresses each conditioned readout relative to the enzyme’s uniform residue-value average to expose chemically conditioned differences.

EnzRef achieves state-of-the-art source prediction on CatPred-DB and zero-shot transfer to the pair-unseen and enzyme-unseen subsets of TurNuP-DB. We evaluate each neural predictor and ablation variant across ten independent training runs with different random seeds. On unseen enzymes, EnzRef achieves a mean *R*^2^ of 0.310 versus 0.259 for the strongest baseline MCKcat, a 19.8% relative improvement. Controlled ablations show that separate role conditioning improves turnover prediction and semantic recomposition further improves activity discrimination. Compared with EnzRef w/o RefCA, the full model reduces heldout log-activity difference RMSE by 6.47% for reactions sharing an enzyme and 5.79% for enzymes sharing a reaction. EnzRef also improves over paired protein-only predictors across six PLMs. With ESM2-650M frozen, it trains 1.72M parameters and achieves a 3.14*×* inference speedup over MCKcat with cached protein and chemical features.

Our contributions are:

- We introduce EnzRef, a lightweight PLM adaptation framework that places cross-modal interaction between chemical semantic decomposition and recomposition. Using frozen PLM features, it models chemical role relations through separately conditioned enzyme responses for turnover prediction.
- We develop RefCA to express chemically conditioned enzyme responses as deviations from an enzyme-specific reference. Centering on the uniform residue-value average exposes conditional differences for more accurate activity discrimination across enzyme–reaction pairs.
- We demonstrate state-of-the-art source prediction on CatPred-DB and zero-shot transfer to unseen pairs and enzymes in TurNuP-DB. EnzRef improves activity-difference prediction and consistently outperforms paired protein-only predictors across six frozen PLMs.

## 2. Related Work

### Enzyme kinetic prediction

Enzyme kinetic predictors differ in how they combine protein and chemical information. UniKP [16] and CataPro [17] combine pretrained protein and molecular features, while CatPred [18] covers complete reactant sets and TurNuP [19] aggregates enzyme and full-reaction predictions. Katalyst [26] aligns sequence, functional text, substrate, and product features before fusion. Chemically conditioned interactions make enzyme features depend on the molecular input: DLKcat [20] uses substrate-guided attention, while PMAK [21] and MCK-cat [22] condition residue aggregation on joint RXNFP embeddings [23]. ERBA [27] further stages conditioning through substrate recognition and pocket geometry. Our En-zRef makes the enzyme’s responses to reactants and products explicit through separate conditioning, then centers and recomposes these responses for turnover prediction.

### Multimodal enzyme–reaction modeling

Cross-modal attention lets one modality condition the features read from another [28], while multimodal factorization separates shared and modality-specific information [29, 30]. In enzyme–chemical modeling, multimodal representations support substrate specificity prediction [31, 32] and enzyme–reaction matching [33]. Reaction encoders organize reactant–product relations through joint graphs [34, 35], and EasIFA [36] incorporates product information into substrate features for active-site annotation. Our EnzRef framework decomposes roles within the chemical modality before enzyme interaction and recomposes their referenced responses afterward. This organization expresses the relation between chemical roles through their separately conditioned enzyme responses.

### Centered and differential attention

Repeated self-attention aggregation can homogenize token representations and cause rank collapse [37, 38]. Centered attention and frequency-aware reweighting mitigate these effects [39, 40], while differential attention suppresses irrelevant-context noise by subtracting learned attention maps [41]. Our RefCA module in EnzRef addresses the mixture of enzyme background and chemically conditioned changes in cross-modal readouts. It measures each response relative to the same enzyme’s uniform residue-value average, exposing conditional deviations for semantic recomposition and cross-sample activity discrimination.

## 3. Method

Given an enzyme sequence *E* and a reaction ℛ, EnzRef predicts *y* = log_10_(*k*_cat_) through semantic decomposition, cross-modal conditioning, and recomposition (Fig. 2). Chemical Semantic Decomposition (CSD) organizes reaction components into reactant and product sets (*C*_*R*_, *C* _*P*_), which separately condition frozen PLM residue features. Reference-Centered Cross-Attention (RefCA) centers each conditioned readout on the enzyme’s uniform residue-value average. Symmetry-Aware Semantic Recomposition (SASR) separately encodes the mean deviation and signed role contrast of these referenced responses to form the task embedding.

**Figure 2.**
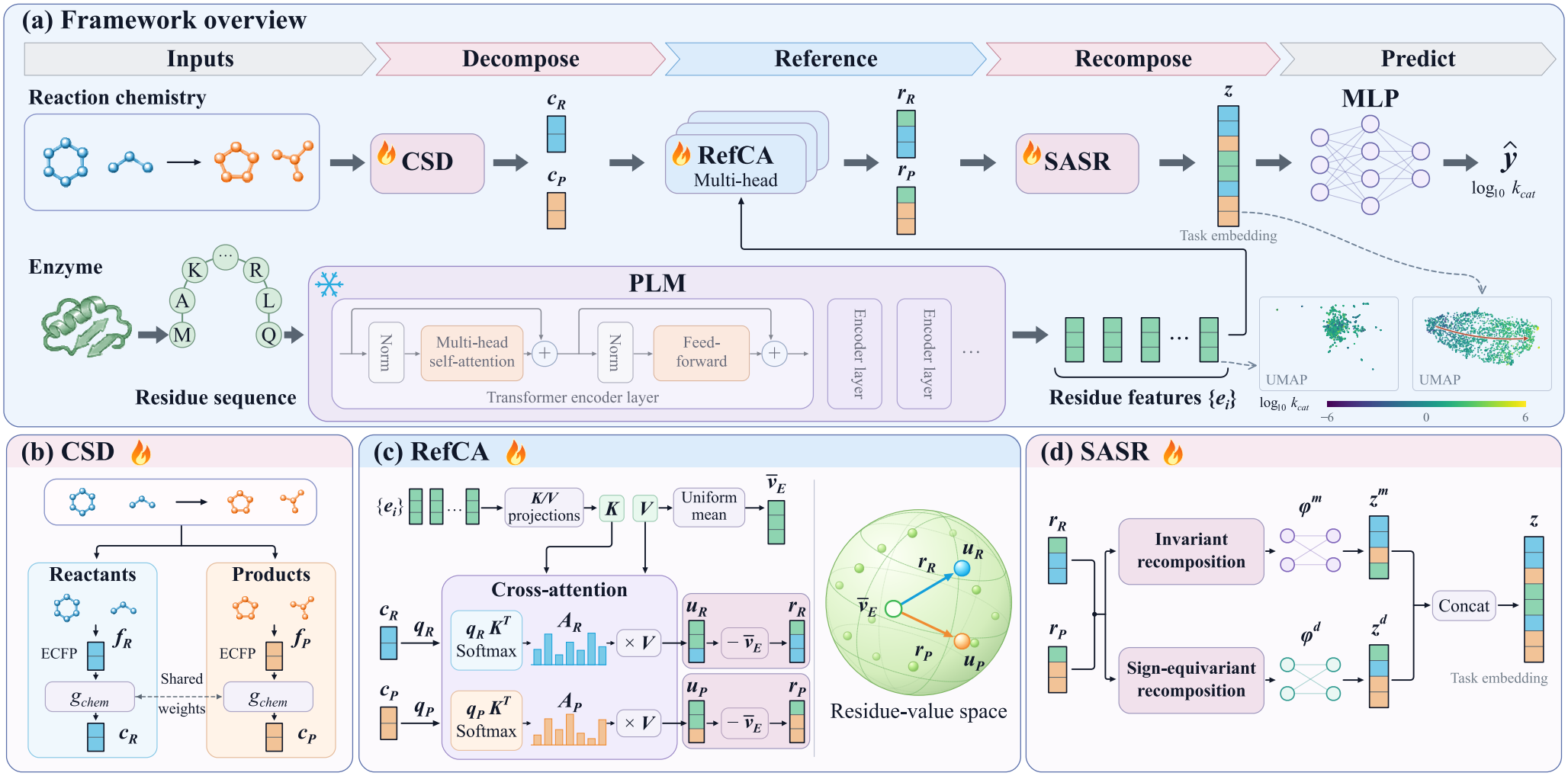
EnzRef decomposes reaction chemistry and recomposes enzyme-referenced responses for activity prediction. (a) Overall pipeline with a frozen PLM. (b) CSD encodes reactant and product conditions separately. (c) RefCA centers cross-attention readouts on the enzyme’s uniform residue-value average. (d) SASR encodes the role-invariant mean deviation and sign-equivariant contrast in separate branches to form a task embedding. Insets show UMAPs of mean-pooled protein and task features, colored by true activity.

### 3.1. Chemical Semantic Decomposition

CSD decomposes the chemical modality according to the reactant and product roles specified by the reaction. Each side supplies a separate condition for enzyme interaction, allowing its role-specific response to be formed before semantic recomposition. For side *s ∈ {R, P}*, we pool the extended-connectivity fingerprints (ECFPs) [42] of its molecular components by bitwise OR and apply a shared chemical encoder *g*_chem_:

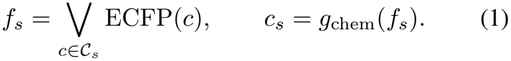

The shared encoder provides a common chemical mapping, while the ordered *c*_*R*_ and *c*_*P*_ channels retain the two roles for separate enzyme conditioning.

### 3.2. Reference-Centered Cross-Attention

Separate chemical queries produce a conditioned enzyme response for each role, but the resulting cross-attention readouts also contain an enzyme-specific residue background. Each readout can be expressed as the enzyme’s uniform residue-value average plus a query-dependent deviation. RefCA centers the readout on this uniform reference to expose the conditional component for learning activity differences across enzyme–reaction pairs.

Let *e*_*i*_ be the frozen PLM representation of residue *i* in an enzyme with *L* valid residues. Learned projections produce residue keys *k*_*i*_ = *g*_*K*_(*e*_*i*_) and values *v*_*i*_ = *g*_*V*_ (*e*_*i*_). Each of *H* heads constructs a query from chemical condition *c*_*s*_ and applies scaled dot-product attention [43]:

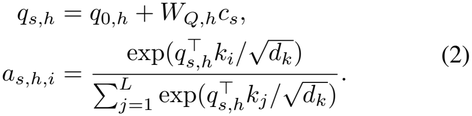

Here, *q*_0,*h*_ is a learned base query and *d*_*k*_ is the key dimension. The query parameters are shared between roles within each head, while all roles and heads use the same keys and values. The resulting readout is 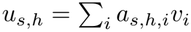.

The learned residue values span an enzyme-specific linear manifold, ℳ_*E*_ = span *{v*_1_, …, *v*_*L*_*}*. Their uniform average provides a query-independent reference shared across roles and heads. We center each conditioned readout on this reference:

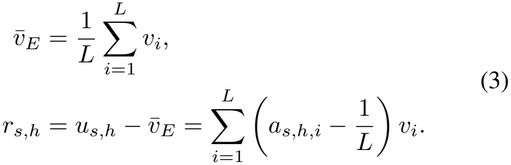

The centered coefficients specify which residue features the query emphasizes or suppresses relative to uniform reading. They sum to zero, so uniform attention produces a zero response; for fixed attention weights, adding a common offset to every value vector leaves *r*_*s,h*_ unchanged. This parameter-free operation supplies the downstream encoders with deviations from each enzyme’s own background for learning activity differences across enzyme–reaction pairs.

### 3.3. Symmetry-Aware Semantic Recomposition

SASR makes the relation between chemical roles explicit through the enzyme-referenced responses produced by RefCA. It first forms their mean deviation and signed role contrast, which separate nonlinear branches encode into a task embedding. For each head, we form

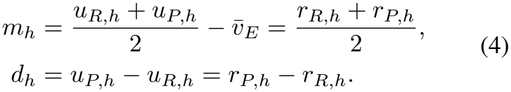

Both recomposed components remain in ℳ_*E*_. The shared reference shifts the mean and cancels from the contrast: *m*_*h*_ represents the average conditional deviation from the enzyme background, while *d*_*h*_ expresses the product-conditioned response relative to the reactant-conditioned response.

Exchanging the two roles maps (*m*_*h*_, *d*_*h*_) to (*m*_*h*_, *™ d*_*h*_). The mean is invariant, whereas the signed role contrast is sign-equivariant. The mean-and-contrast representation preserves both input responses, since *r*_*R,h*_ = *m*_*h*_ *™ d*_*h*_*/*2 and *r*_*P,h*_ = *m*_*h*_ + *d*_*h*_*/*2. We use this role-exchange structure to organize the inputs to separate nonlinear branches:

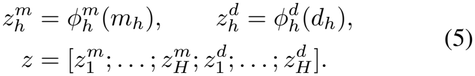

Each *ϕ* consists of layer normalization, an affine projection, and GELU, with independent parameters for each head and relation type; [;] denotes concatenation.

### 3.4. Prediction and Learning

A multilayer perceptron maps *z* to a standardized log-activity prediction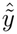. Let 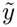 denote the label standardized using the source-training mean and standard deviation. We minimize

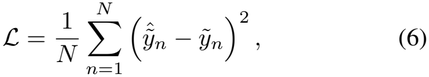

where *N* is the number of training records. Optimization updates the chemical encoder, RefCA projections and queries, SASR branches, and prediction head while keeping the PLM frozen. At inference, predictions are restored to the log_10_(*k*_cat_) scale using the same training statistics.

## 4. Experiments

We first evaluate source-domain prediction, zero-shot transfer, and adaptation across frozen PLMs, followed by activity discrimination and representation analysis. We then test decomposition before enzyme interaction and recomposition afterward, isolate the contribution of reference centering, and assess inference efficiency.

### 4.1. Experimental Setup

#### Data and transfer protocol

We preserved the original training, validation, and test assignments of CatPred-DB [18] and removed records with missing products, con-firmed product–label mismatches, or unparsable molecular structures. We identified a unit-conversion error in CatPred-DB’s released preprocessing of SABIO-RK [44] records and corrected 1,918 affected labels using the source measurements. The resulting dataset contains 16,587 training, 1,837 validation, and 2,049 held-out records. The target is *y* = log_10_(*k*_cat_), with *k*_cat_ expressed in s^*™*1^. For zero-shot transfer, we used TurNuP-DB records [19] with complete reaction inputs. The pair-unseen subset contains 2,746 records whose combinations of enzyme sequence and reactants are absent from source training and validation. The nested enzyme-unseen subset contains 1,008 records whose exact enzyme sequence is absent from both source splits. We applied fixed source-trained models without target-domain training or tuning. Appendix A details data curation and external subset construction.

#### Baselines

We compared against DLKcat [20], CatPred [18], TurNuP [19], UniKP [16], CataPro [17], PMAK [21], and MCKcat [22], starting from their released implementations and configurations. For a fair comparison, PLM-based methods used the same frozen ESM2-650M enzyme encoder [7]. EnzRef and neural baselines used a common five-candidate learning-rate grid and were trained on the source training split, with learning rates and checkpoints selected by validation RMSE. Tree-based UniKP and TurNuP were fitted once on the combined training and validation records. Appendix B provides implementation and evaluation details.

#### Metrics and repeated training

We report *R*^2^, MAE, and Spearman rank correlation (SRCC) for prediction accuracy and activity ordering in log_10_(*k*_cat_) space. For neural methods, we report ten-run means *±* standard deviations and ensemble scores from equally averaged log-space predictions. Complete results, including RMSE and Pearson correlation (PCC), appear in Appendix J.

#### Implementation

We trained EnzRef with AdamW [45], a learning rate of 10^*™*3^, and batch size 512. Architecture and optimization details appear in Appendix C.

### 4.2. Source-Domain Prediction and Zero-Shot Transfer

#### Source-domain prediction

EnzRef achieves state-of-the-art source prediction and overall rank agreement across all three metrics in both evaluation modes (Table 1). Relative to MCKcat, EnzRef improves mean single-model *R*^2^ from 0.553 to 0.614 and SRCC from 0.732 to 0.758. Its mean single-model *R*^2^ also exceeds MCKcat’s ensemble score of 0.599. These gains show that EnzRef improves both turnover estimation and activity ordering through lightweight multimodal adaptation of frozen protein features.

**Table 1.**
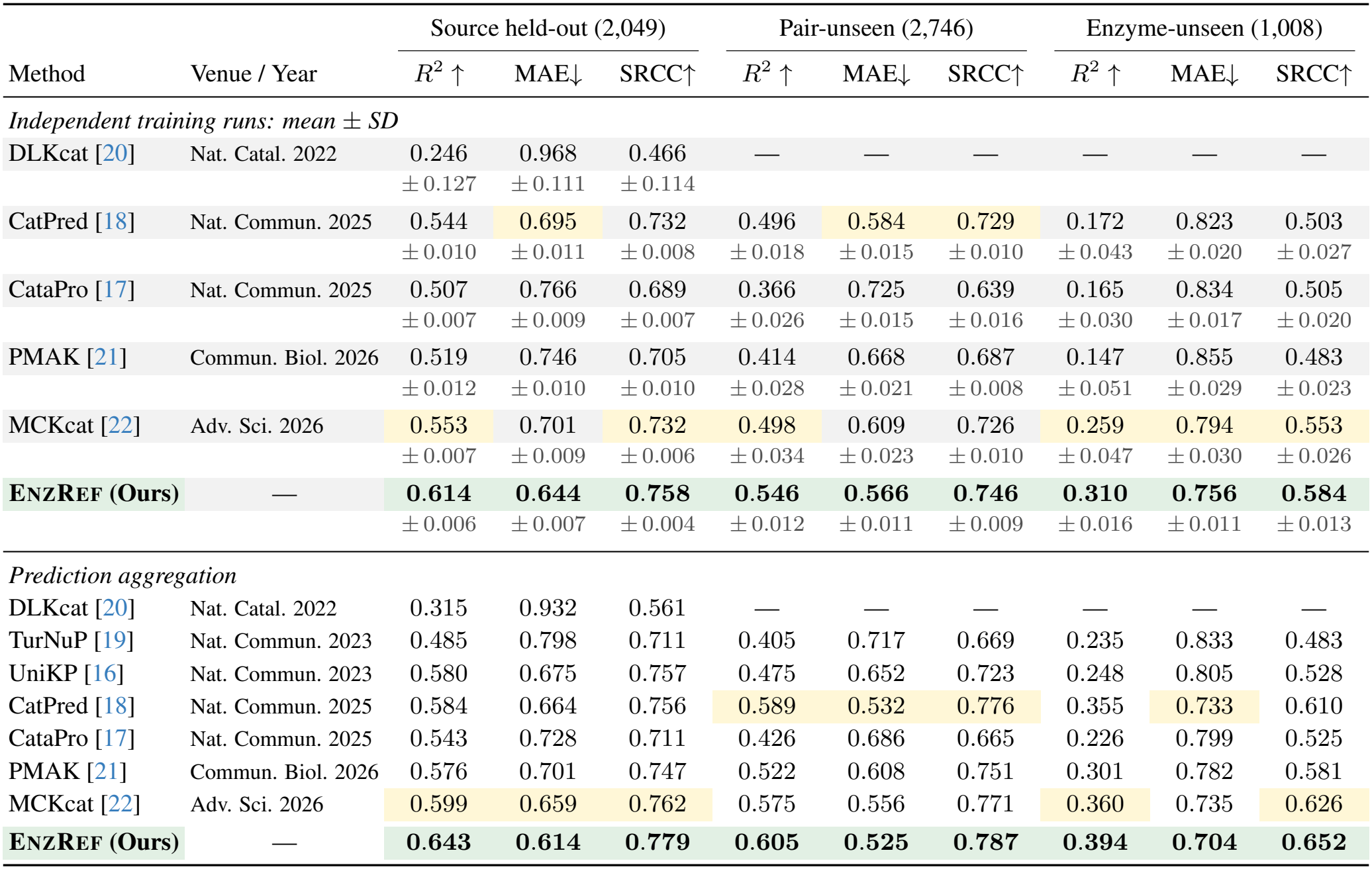
Source prediction and zero-shot transfer. Upper block: ten-run means (gray rows) with *±* population SD below; lower block: aggregated predictions. UniKP and TurNuP use internal aggregation. CatPred uses 2,048 source records; DLKcat’s external scores are omitted for incomplete coverage (Appendix B). **Green/bold** and yellow mark the best and second-best values per block before rounding.

| Method | Venue / Year | Source held-out (2,049) |  |  | Pair-unseen (2,746) |  |  | Enzyme-unseen (1,008) |  |  |
|---|---|---|---|---|---|---|---|---|---|---|
|  |  | *R*<sup>2</sup> ↑ | MAE↓ | SRCC↑ | *R*<sup>2</sup> ↑ | MAE↓ | SRCC↑ | *R*<sup>2</sup> ↑ | MAE↓ | SRCC↑ |
| *Independent training runs: mean ± SD* |  |  |  |  |  |  |  |  |  |  |
| DLKcat [20] | Nat. Catal. 2022 | 0.246<br>±0.127 | 0.968<br>±0.111 | 0.466<br>±0.114 | — | — | — | — | — | — |
| CatPred [18] | Nat. Commun. 2025 | 0.544<br>±0.010 | 0.695<br>±0.011 | 0.732<br>±0.008 | 0.496<br>±0.018 | 0.584<br>±0.015 | 0.729<br>±0.010 | 0.172<br>±0.043 | 0.823<br>±0.020 | 0.503<br>±0.027 |
| CataPro [17] | Nat. Commun. 2025 | 0.507<br>±0.007 | 0.766<br>±0.009 | 0.689<br>±0.007 | 0.366<br>±0.026 | 0.725<br>±0.015 | 0.639<br>±0.016 | 0.165<br>±0.030 | 0.834<br>±0.017 | 0.505<br>±0.020 |
| PMAK [21] | Commun. Biol. 2026 | 0.519<br>±0.012 | 0.746<br>±0.010 | 0.705<br>±0.010 | 0.414<br>±0.028 | 0.668<br>±0.021 | 0.687<br>±0.008 | 0.147<br>±0.051 | 0.855<br>±0.029 | 0.483<br>±0.023 |
| MCKcat [22] | Adv. Sci. 2026 | 0.553<br>±0.007 | 0.701<br>±0.009 | 0.732<br>±0.006 | 0.498<br>±0.034 | 0.609<br>±0.023 | 0.726<br>±0.010 | 0.259<br>±0.047 | 0.794<br>±0.030 | 0.553<br>±0.026 |
| **EnzRef (Ours)** | — | **0.614**<br>±0.006 | **0.644**<br>±0.007 | **0.758**<br>±0.004 | **0.546**<br>±0.012 | **0.566**<br>±0.011 | **0.746**<br>±0.009 | **0.310**<br>±0.016 | **0.756**<br>±0.011 | **0.584**<br>±0.013 |
| *Prediction aggregation* |  |  |  |  |  |  |  |  |  |  |
| DLKcat [20] | Nat. Catal. 2022 | 0.315 | 0.932 | 0.561 | — | — | — | — | — | — |
| TurNuP [19] | Nat. Commun. 2023 | 0.485 | 0.798 | 0.711 | 0.405 | 0.717 | 0.669 | 0.235 | 0.833 | 0.483 |
| UniKP [16] | Nat. Commun. 2023 | 0.580 | 0.675 | 0.757 | 0.475 | 0.652 | 0.723 | 0.248 | 0.805 | 0.528 |
| CatPred [18] | Nat. Commun. 2025 | 0.584 | 0.664 | 0.756 | 0.589 | 0.532 | 0.776 | 0.355 | 0.733 | 0.610 |
| CataPro [17] | Nat. Commun. 2025 | 0.543 | 0.728 | 0.711 | 0.426 | 0.686 | 0.665 | 0.226 | 0.799 | 0.525 |
| PMAK [21] | Commun. Biol. 2026 | 0.576 | 0.701 | 0.747 | 0.522 | 0.608 | 0.751 | 0.301 | 0.782 | 0.581 |
| MCKcat [22] | Adv. Sci. 2026 | 0.599 | 0.659 | 0.762 | 0.575 | 0.556 | 0.771 | 0.360 | 0.735 | 0.626 |
| **EnzRef (Ours)** | — | **0.643** | **0.614** | **0.779** | **0.605** | **0.525** | **0.787** | **0.394** | **0.704** | **0.652** |

#### Zero-shot transfer

EnzRef leads all three metrics on unseen pairs and enzymes in both evaluation modes. With fixed source-trained models, mean single-model *R*^2^ reaches 0.546 on unseen pairs versus 0.498 for MCKcat, a 9.7% relative improvement. On unseen enzymes, *R*^2^ reaches 0.310, a 19.8% relative improvement over MCKcat, while mean SRCC rises from 0.553 to 0.584. These results support reusing source-trained models to estimate and order candidate activities before target-specific measurements are available.

### 4.3. Multimodal Adaptation across PLMs

EnzRef improves ensemble *R*^2^ and SRCC over paired protein-only predictors across five ESM2 sizes [7] and ProtT5-XL [8] on the source held-out set and both external subsets (Fig. 3). Each pair uses the same frozen encoder, with mean-pooled residue features for the protein-only predictor. Downstream modules were trained separately for each PLM using the default ESM2-650M training hyperparameters (Appendix D). Relative gains range from 16.6–20.7% in source *R*^2^ and 12.8–24.5% in enzyme-unseen SRCC.

**Figure 3.**
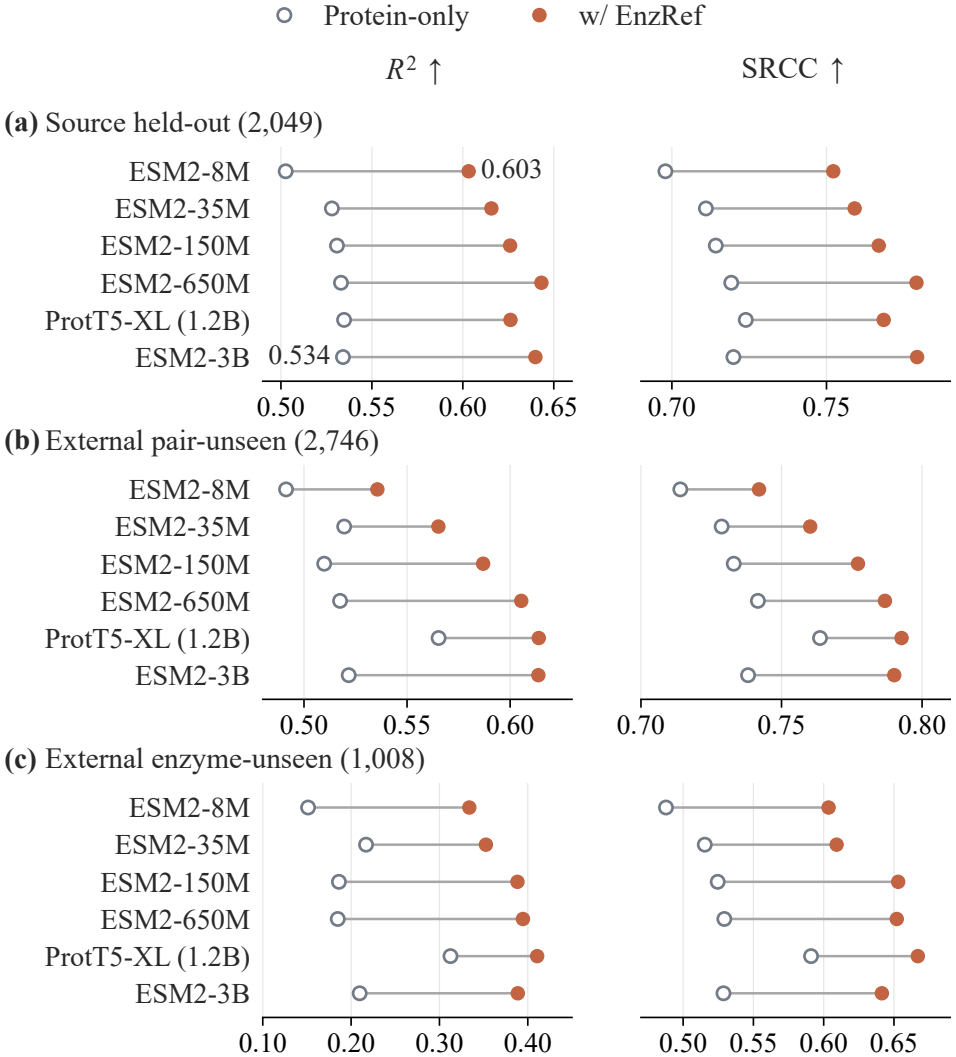
EnzRef improves prediction across frozen PLMs. Ten-model ensemble *R*^2^ and SRCC. Lines connect protein-only and EnzRef results for the same PLM. Full metrics appear in Appendix J.

Reaction-conditioned adaptation enables a compact PLM to outperform a much larger protein-only predictor. On the source held-out set, EnzRef with ESM2-8M achieves an ensemble *R*^2^ of 0.603, compared with 0.534 for the ESM2-3B protein-only predictor (Fig. 3a). Mean single-model *R*^2^ also favors EnzRef, at 0.561 versus 0.515. These results illustrate the value of adapting protein features to reaction chemistry: increasing encoder size alone does not supply the chemical context needed to distinguish activities across reactions of the same enzyme.

### 4.4. Activity Discrimination and Representation Analysis

Beyond overall rank agreement, we test whether EnzRef captures activity differences between pairs sharing an enzyme or a reaction, then examine how activity is organized in its learned representations.

#### Discriminating conditional activity differences

We measure activity discrimination in two held-out settings: different reactions catalyzed by the same exact enzyme sequence, and different enzymes catalyzing the same canonical reaction. Within each eligible group, duplicate enzyme– reaction records were averaged before enumerating all unordered pairs. For *G* groups with pair sets *P*_*g*_, we computed

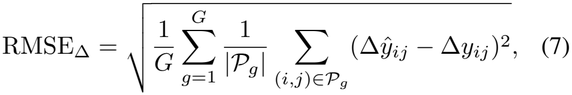

where Δ*y*_*ij*_ = *y*_*j*_ *™ y*_*i*_ and Δ*ŷ*_*ij*_ = *ŷ*_*j*_ *™ ŷ*_*i*_. Equal group weighting prevents enzymes or reactions with many records from dominating the metric.

EnzRef predicts activity differences more accurately than PMAK and MCKcat when either the enzyme or the reaction is held fixed (Table 2). Compared with PMAK, the strongest baseline on both tasks, mean difference RMSE falls by 5.63% for shared enzymes and 6.18% for shared reactions. Appendix E details grouping and pair counts.

**Table 2.**
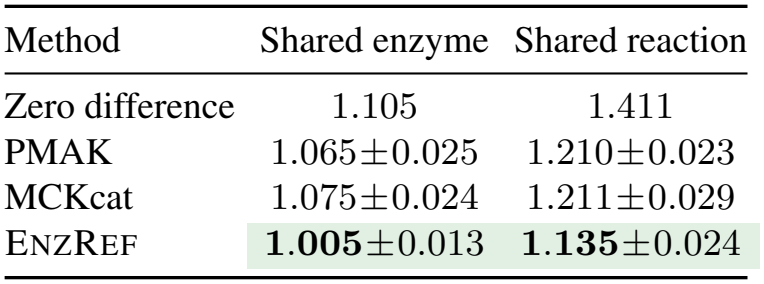
More accurate prediction of activity differences. Difference RMSE on held-out groups: 317 enzymes (864 pairs) and 190 reactions (458 pairs). Values are means *±* population SD over ten training runs; lower is better. The zero-difference predictor assigns identical activity within a group. **Green/bold** marks the lowest mean in each column.

| Method | Shared enzyme | Shared reaction |
|---|---|---|
| Zero difference | 1.105 | 1.411 |
| PMAK | 1.065±0.025 | 1.210±0.023 |
| MCKcat | 1.075±0.024 | 1.211±0.029 |
| ENZREF | **1.005**±0.013 | **1.135**±0.024 |

#### Held-out representation analysis

EnzRef reduces the strong alignment between sample representations seen in both ESM-mean and MCKcat (Fig. 4b). In the original feature spaces, mean pairwise cosine similarity is 0.181 for EnzRef, compared with 0.900 for ESM-mean and 0.889 for MCKcat. Its broader cosine distribution reflects more varied feature directions across enzyme–reaction pairs.

**Figure 4.**
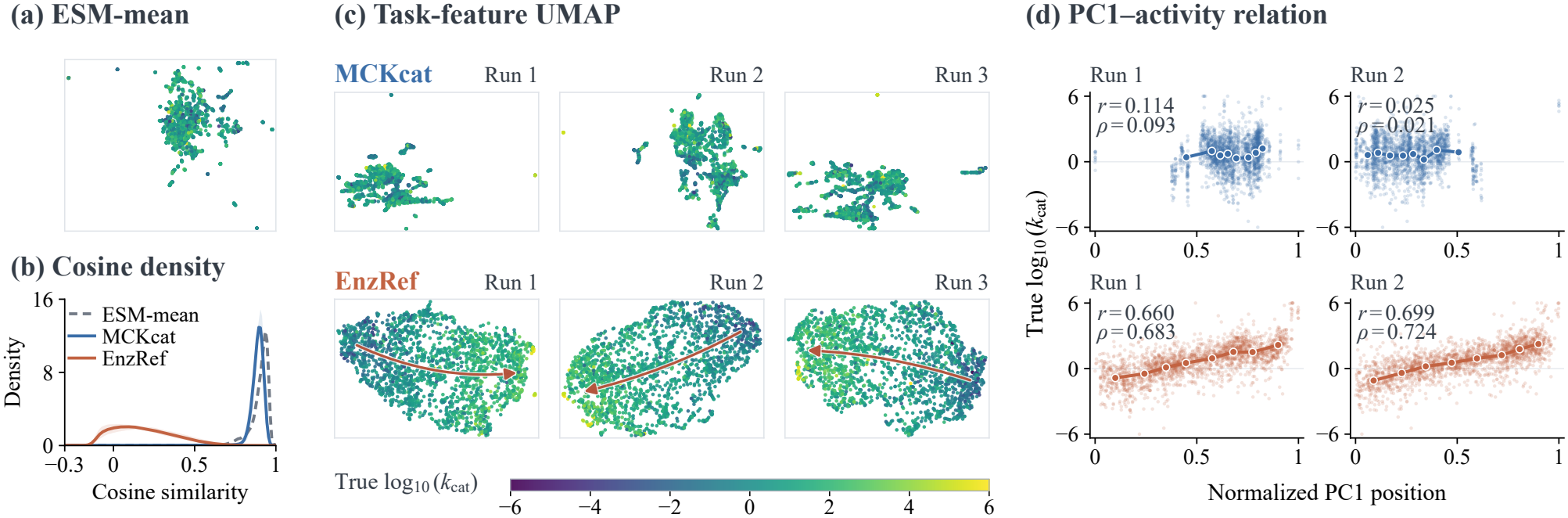
EnzRef features show clearer activity organization. (a,c) UMAPs fitted from features and colored by true log-activity; matching run numbers indicate shared training seeds. (b) Original-space cosine densities: ten-run means and min–max bands for task models. (d) Activity versus normalized UMAP PC1 position; lines connect eight group means, and *r*/*ρ* denote sample-level PCC/SRCC. See Appendix G for label-based orientation and UMAP guides.

This reduced alignment is accompanied by clearer activity gradients in EnzRef’s UMAPs (Fig. 4c,d). For the first two matched runs, sample-level PCC between UMAP PC1 position and true log-activity reaches 0.660 and 0.699 for EnzRef. The corresponding MCKcat values are 0.114 and 0.025. Together with the lower conditional-difference errors, these observations support EnzRef’s ability to preserve activity-relevant differences between enzyme–reaction pairs. All-run projections and representation analyses appear in Appendix F.

### 4.5. Ablation Studies

We first test the benefit of separating chemical roles before enzyme interaction and recomposing their responses afterward, then isolate the contribution of enzyme-background referencing. All variants use frozen ESM2-650M features and the same ten training seeds; Tables 3 and 4 report source held-out single-model means *±* SD. Complete metrics appear in Appendix J.

**Table 3.**
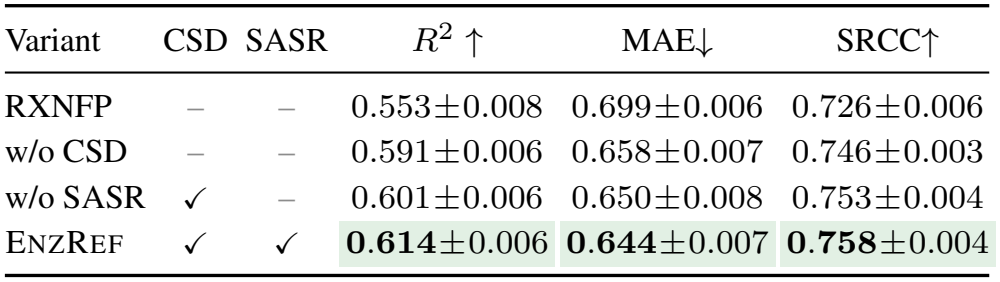
Chemical semantic decomposition and recomposition improve prediction and ordering. Checks indicate enabled modules; all variants retain RefCA. Source held-out results are single-model means *±* population SD over ten runs with frozen ESM2-650M. **Green/bold** marks the best mean in each metric column.

| Variant | CSD | SASR | *R*<sup>2</sup> ↑ | MAE↓ | SRCC↑ |
|---|---|---|---|---|---|
| RXNFP | – | – | 0.553±0.008 | 0.699±0.006 | 0.726±0.006 |
| w/o CSD | – | – | 0.591±0.006 | 0.658±0.007 | 0.746±0.003 |
| w/o SASR | ✓ | – | 0.601±0.006 | 0.650±0.008 | 0.753±0.004 |
| ENZREF | ✓ | ✓ | **0.614**±0.006 | **0.644**±0.007 | **0.758**±0.004 |

**Table 4.**
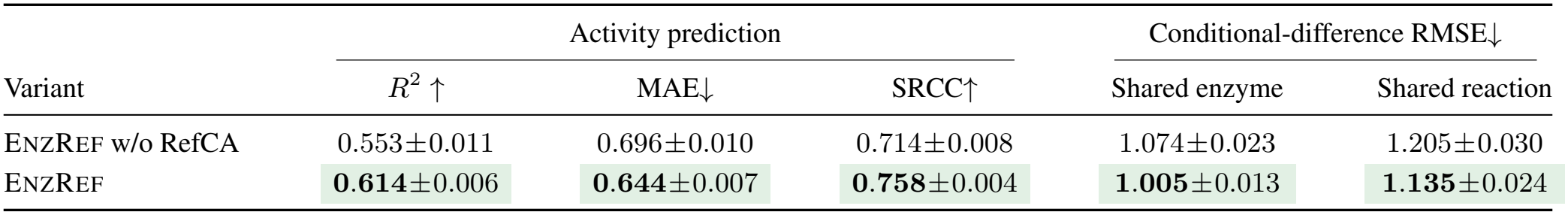
RefCA improves prediction and conditional activity discrimination. EnzRef w/o RefCA uses uncentered cross-attention readouts, retaining CSD, SASR, and the same parameter count. Entries are source held-out means *±* population SD over ten matched runs. **Green/bold** marks the better mean.

|  | Activity prediction |  |  | Conditional-difference RMSE↓ |  |
|---|---|---|---|---|---|
| Variant | *R*<sup>2</sup> ↑ | MAE↓ | SRCC↑ | Shared enzyme | Shared reaction |
| EnzRef w/o RefCA | 0.553±0.011 | 0.696±0.010 | 0.714±0.008 | 1.074±0.023 | 1.205±0.030 |
| EnzRef | **0.614**±0.006 | **0.644**±0.007 | **0.758**±0.004 | **1.005**±0.013 | **1.135**±0.024 |

#### Chemical Semantic Decomposition and Recomposition

Conditioning the enzyme separately on reactants and products improves activity prediction and ordering (Table 3). RXNFP uses a joint reaction embedding, while w/o CSD concatenates the two sides’ ECFPs in fixed order; both provide a single condition for enzyme interaction. With the same reactant and product ECFPs, w/o SASR forms separate enzyme responses for the two roles and encodes them independently. Relative to w/o CSD, this raises mean *R*^2^ from 0.591 to 0.601 and SRCC from 0.746 to 0.753.

SASR further improves activity discrimination by recomposing these responses according to their chemical role relation. With the same parameter count, encoding their mean deviation and signed role contrast instead of the two sides separately raises mean *R*^2^ from 0.601 to 0.614 and SRCC from 0.753 to 0.758. Conditional-difference RMSE also decreases from 1.023 to 1.005 for shared enzymes and from 1.162 to 1.135 for shared reactions (Appendix H.1).

Together, these comparisons show that separating chemical roles before enzyme interaction and recomposing their responses afterward improves activity prediction and discrimination.

#### Reference-Centered Cross-Attention

RefCA improves both turnover prediction and activity discrimination across enzyme–reaction pairs (Table 4). Replacing uncentered cross-attention readouts with enzyme-referenced deviations raises mean *R*^2^ from 0.553 to 0.614. Conditional-difference RMSE decreases by 6.47% for reactions sharing an enzyme and 5.79% for enzymes sharing a reaction. Both errors decrease in all ten paired runs, supporting more accurate discrimination of activity differences even when samples share one of their inputs.

Reference centering reduces mean cross-sample readout cosine from 0.429 to 0.073 (Fig. 5) and within-sample reactant–product cosine from 0.961 to 0.649 (Fig. 6). Together with lower conditional-difference errors, these changes support learning activity differences from more directionally distinct chemical responses relative to each enzyme’s residue background.

**Figure 5.**
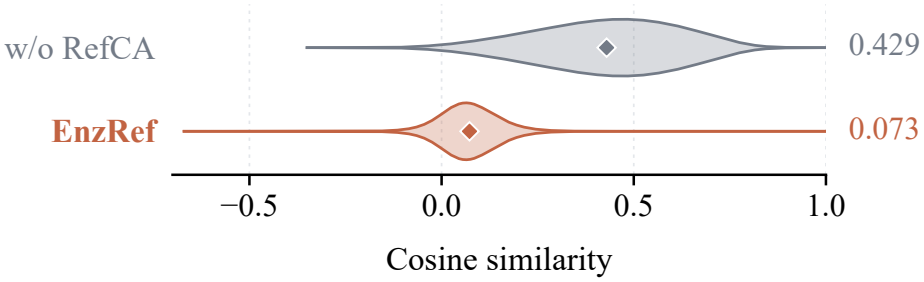
RefCA reduces cross-sample readout alignment. Cosine of concatenated reactant/product readouts before nonlinear encoding. Violins show ten-run mean training-set densities; diamonds mark mean cosines.

**Figure 6.**
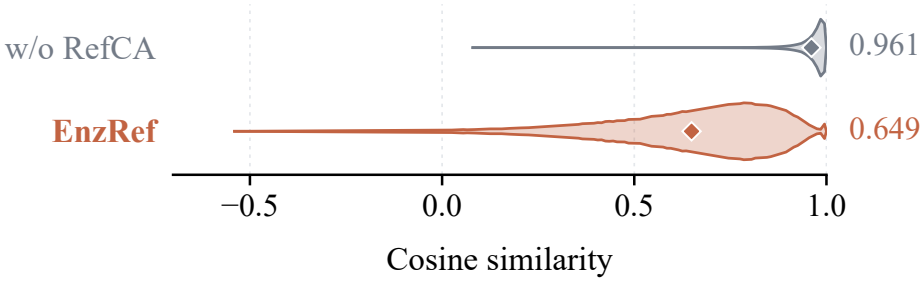
RefCA reduces alignment between chemical roles. Within-sample reactant–product cosine before nonlinear encoding. Density aggregation and mean markers follow Fig. 5.

### 4.6 Inference Efficiency

Among the PLM-based methods in Fig. 7, EnzRef achieves the highest source *R*^2^ with the fewest parameters beyond the shared frozen encoder (1.72M). With cached inputs, held-out inference takes 0.775 *±* 0.035 seconds per checkpoint, corresponding to speedups of 3.14*×* over MCKcat and 9.84*×* over CatPred. This efficiency supports highthroughput virtual screening to prioritize enzyme–reaction pairs by predicted activity.

**Figure 7.**
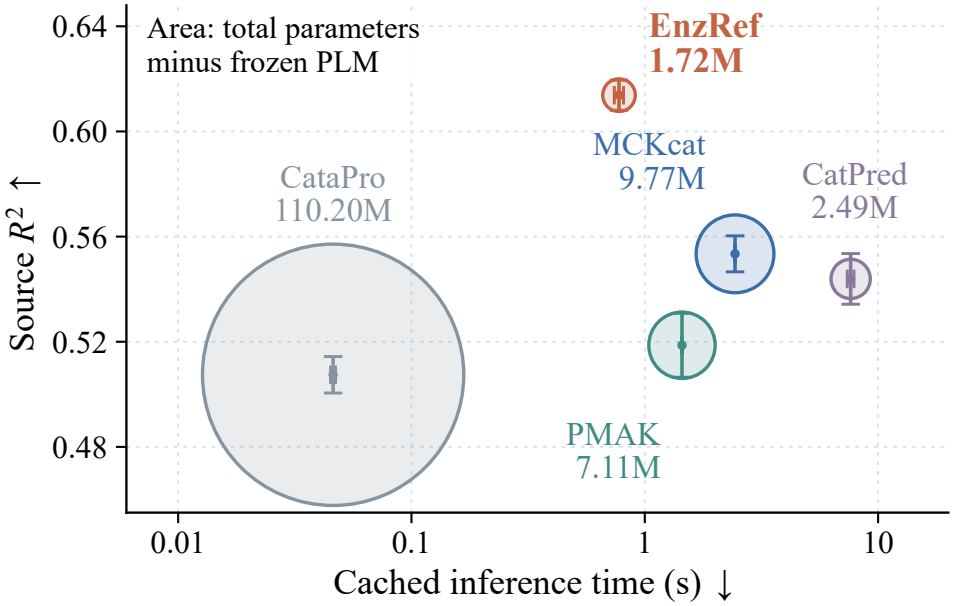
Accurate prediction at low inference cost. Source *R*^2^ versus per-checkpoint runtime with cached inputs (logarithmic time axis). Bubble area denotes parameters excluding the frozen PLM; error bars show ten-run SD. Details appear in Appendix I.

## 5. Conclusion

We presented EnzRef, a lightweight multimodal framework that adapts frozen PLM features for enzyme turnover prediction. EnzRef separates chemical roles before enzyme interaction and recomposes their responses afterward to make role relations explicit. RefCA expresses each conditioned response as a deviation from the enzyme’s own residue background for learning activity differences. Experiments demonstrate state-of-the-art source prediction on CatPred-DB and zero-shot transfer to TurNuP-DB, with consistent adaptation gains across six PLMs. EnzRef also predicts activity differences more accurately between pairs sharing an enzyme or a reaction. With its predictive accuracy, lightweight adaptation, and low inference cost, EnzRef supports high-throughput virtual screening of enzyme–reaction pairs with known reactant and product chemistry. Extending the framework to settings where product information is unavailable or incomplete would broaden its applicability.

## Data and Code Availability

The original datasets are available from CatPred-DB [18] and TurNuP [19]. The EnzRef code and processed data will be made publicly available upon publication of the peerreviewed article.

## LLM Usage Statement

During the preparation of this work, the authors used Chat-GPT to assist with manuscript organization, language editing and refinement, and the development of visual concepts and layouts for illustrative figures. All AI-assisted text and figure designs were reviewed, revised, and approved by the authors. The authors take full responsibility for the entire work, including its scientific content, experimental results, and figures.

## Supplementary Information

### A. Data Preparation and External Evaluation

#### A.1. Source Data Curation

##### Filtering while preserving the original split

We started from the released CatPred-DB random split [18] and retained each record’s original training, validation, or test assignment. We excluded records with missing product SMILES, a confirmed mismatch between the assigned product and the source measurement providing the group’s maximum value, or a molecular component that RDKit could not parse. These criteria identify 2,335, 355, and one record, respectively; 13 records satisfy both of the first two criteria. Their union removes 2,678 records, leaving 20,473 records with the split sizes in Table 5.

**Table 5.**
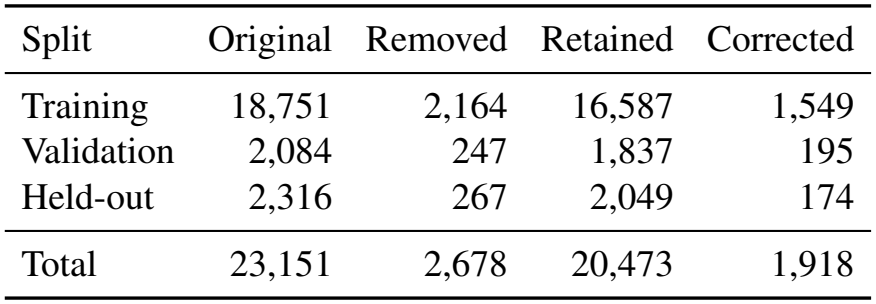
Source data curation preserves the released split assignments. “Corrected” counts label updates among retained records.

| Split | Original | Removed | Retained | Corrected |
|---|---|---|---|---|
| Training | 18,751 | 2,164 | 16,587 | 1,549 |
| Validation | 2,084 | 247 | 1,837 | 195 |
| Held-out | 2,316 | 267 | 2,049 | 174 |
| Total | 23,151 | 2,678 | 20,473 | 1,918 |

##### Unit verification and label correction

The released preprocessing contains incorrect conversions of SABIO-RK turnover measurements [44]. For example, a value of 2.42 min^*™*1^ appears as 145.2 s^*™*1^ in the processed records, whereas the source database gives 2.42*/*60 = 0.04033 s^*™*1^. The processed value multiplies by 60 instead of dividing by 60. The audit also identifies analogous conversion errors for measurements in h^*™*1^.

We reconstructed each CatPred group using the full enzyme sequence and the sorted reactant SMILES components, preserving repeated components as in the original grouping. For each group, we traced candidate measurements from both BRENDA [46] and SABIO-RK, since the maximum can originate from either database. We checked SABIO-RK’s original value and unit against its standardized value, and matched BRENDA records using the available enzyme identifiers, EC number, substrate, and measurement value. We then recomputed the group maximum from the standardized values in s^*™*1^ and took its base-10 logarithm. Recomputing the maximum is necessary because unit correction can change which measurement determines the target.

We applied corrections supported by the frozen source audit, yielding 1,549 changed training labels, 195 validation labels, and 174 held-out labels. This correction changes only the target and its derived label files; it preserves the retained records, chemical inputs, enzyme sequences, and split assignments. Source training and evaluation use this corrected version throughout. The frozen manifests record the source files, filtering rules, split counts, stable record identifiers, and file checksums; separate ledgers retain the exclusions and label changes.

#### A.2. External Records and Overlap Audit

##### External pool

We used the TurNuP data [19] distributed in PMAK’s warm-split export [21]. Pooling the training, validation, and test files of one fold gives 4,271 unique records and 2,800 unique enzyme sequences, with both reaction sides available. These external split names identify the source files only: every record is evaluated by the fixed source-trained models. We retain the external snapshot’s log-space target values and apply no target-domain training or tuning.

##### Nested evaluation subsets

We audited overlap against the union of the corrected source training and validation sets. Enzymes are identified by exact matches of their full amino-acid sequences. For each chemical side, we canonicalize component SMILES with RDKit, remove duplicate components, and sort them, while preserving reactant– product direction. Let *E, R*, and *P* denote the resulting enzyme, reactant-side, and product-side keys. The external pool contains 921 records whose (*E, R, P*) key occurs in the source training or validation set. Removing these leaves 3,350 full-input-unseen records. Excluding another 604 records with a source-seen (*E, R*) key gives the 2,746 pairunseen records used in the main comparison. Requiring *E* itself to be absent gives the nested 1,008-record enzyme-unseen subset.

##### Results on the complete external pool

Table 6 supplements the main comparison with results on all 4,271 external records. EnzRef has the highest *R*^2^ and lowest RMSE and MAE in both evaluation modes among methods covering this pool. Because the pool includes the 921 source-overlapping complete inputs, the main paper uses the pairunseen and enzyme-unseen subsets to assess transfer to unseen inputs.

**Table 6.**
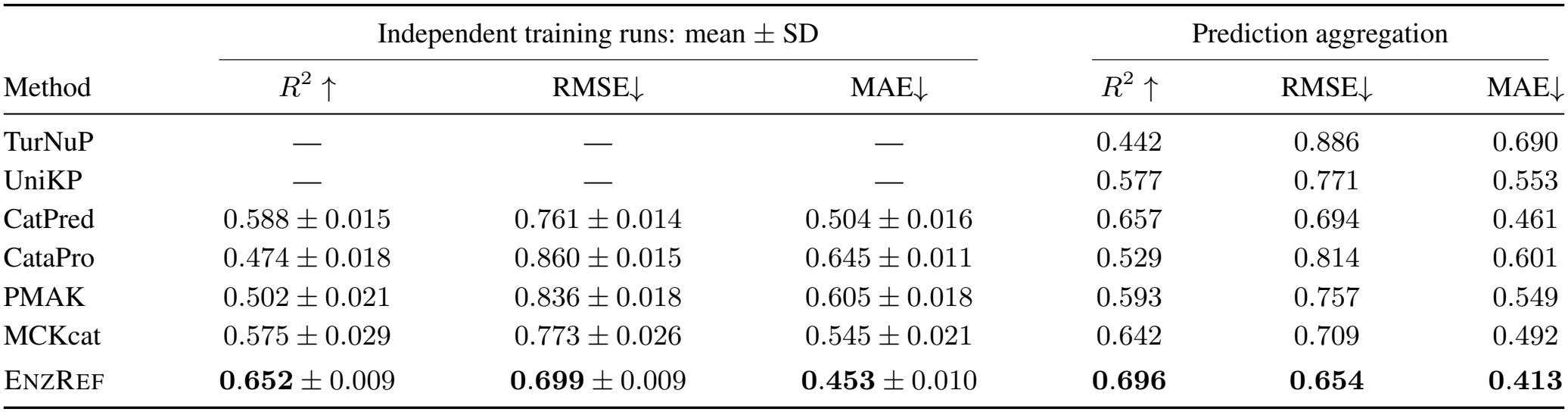
Prediction on the complete TurNuP external pool (4,271 records). All models are fixed after source-domain training. The pool includes 921 complete inputs also present in source training or validation. Neural-model scores are mean *±* population SD over ten runs; prediction aggregation averages their log-space predictions. TurNuP and UniKP appear only under prediction aggregation, following Table 1. DLKcat is excluded from this full-pool comparison because its frozen vocabulary does not cover every record. Bold indicates the best value within each evaluation mode.

| Method | Independent training runs: mean ± SD |  |  | Prediction aggregation |  |  |
|---|---|---|---|---|---|---|
|  | *R*<sup>2</sup> ↑ | RMSE↓ | MAE↓ | *R*<sup>2</sup> ↑ | RMSE↓ | MAE↓ |
| TurNuP | — | — | — | 0.442 | 0.886 | 0.690 |
| UniKP | — | — | — | 0.577 | 0.771 | 0.553 |
| CatPred | 0.588 ± 0.015 | 0.761 ± 0.014 | 0.504 ± 0.016 | 0.657 | 0.694 | 0.461 |
| CataPro | 0.474 ± 0.018 | 0.860 ± 0.015 | 0.645 ± 0.011 | 0.529 | 0.814 | 0.601 |
| PMAK | 0.502 ± 0.021 | 0.836 ± 0.018 | 0.605 ± 0.018 | 0.593 | 0.757 | 0.549 |
| MCKcat | 0.575 ± 0.029 | 0.773 ± 0.026 | 0.545 ± 0.021 | 0.642 | 0.709 | 0.492 |
| ENZREF | **0.652** ± 0.009 | **0.699** ± 0.009 | **0.453** ± 0.010 | **0.696** | **0.654** | **0.413** |

### B. Baseline Implementations and Evaluation

#### Inputs and encoder adaptation

Table 7 summarizes the inputs and released prediction components retained for each baseline. All PLM-based methods use the same frozen ESM2-650M layer-33 features, extracted and stored in FP32. For PMAK, MCKcat, and CataPro, we adjust the protein-input layer dimensions from 1,024 to 1,280; TurNuP and UniKP receive the 1,280-dimensional sequence mean. DLKcat retains its learned sequence encoder. On the source dataset, the single-substrate inputs of DLKcat, CataPro, and UniKP follow the CatPred authors’ released substrate mapping. Each selected SMILES is one complete component of the record’s reactant string.

**Table 7.**
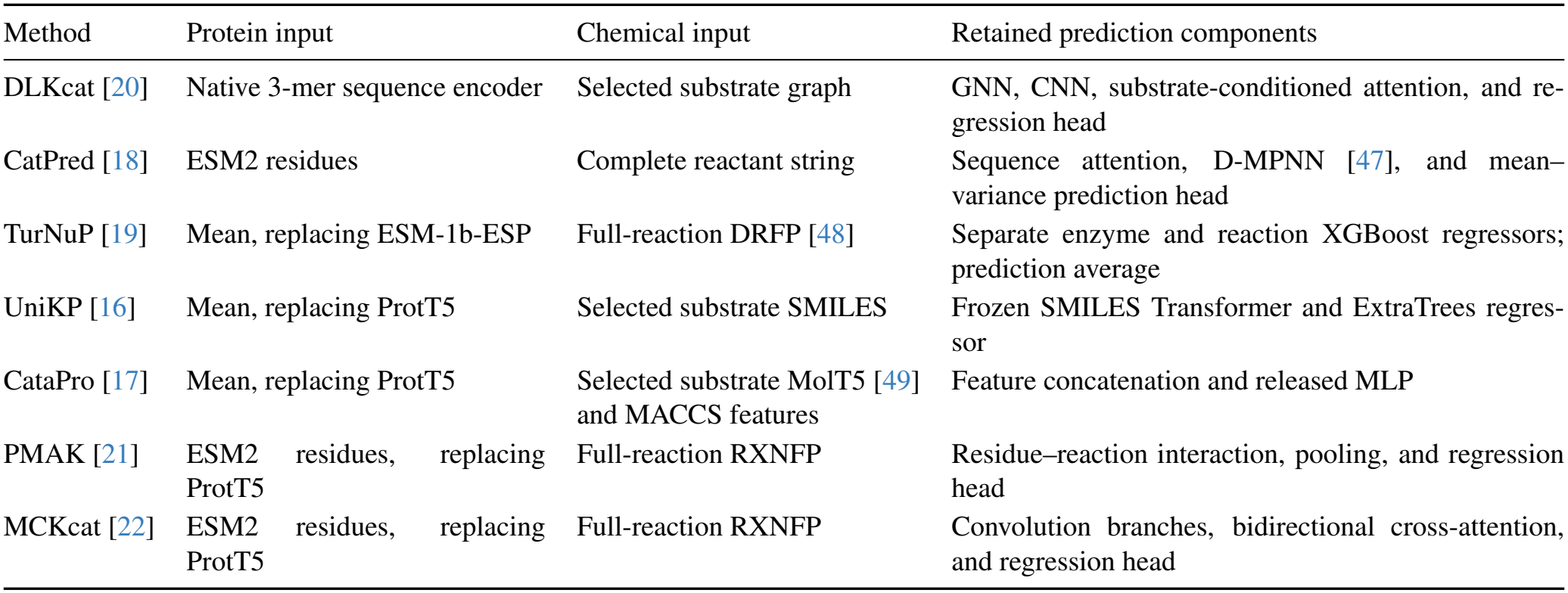
Baseline inputs and retained prediction components. “Mean” denotes the uniform mean of frozen ESM2-650M residue features. The chemical-input column distinguishes a selected substrate, the reactant side, and the complete reaction.

| Method | Protein input | Chemical input | Retained prediction components |
|---|---|---|---|
| DLKcat [20] | Native 3-mer sequence encoder | Selected substrate graph | GNN, CNN, substrate-conditioned attention, and re-gression head |
| CatPred [18] | ESM2 residues | Complete reactant string | Sequence attention, D-MPNN [47], and mean–variance prediction head |
| TurNuP [19] | Mean, replacing ESM-1b-ESP | Full-reaction DRFP [48] | Separate enzyme and reaction XGBoost regressors; prediction average |
| UniKP [16] | Mean, replacing ProtT5 | Selected substrate SMILES | Frozen SMILES Transformer and ExtraTrees regres-sor |
| CataPro [17] | Mean, replacing ProtT5 | Selected substrate MolT5 [49] and MACCS features | Feature concatenation and released MLP |
| PMAK [21] | ESM2 residues, replacing ProtT5 | Full-reaction RXNFP | Residue–reaction interaction, pooling, and regression head |
| MCKcat [22] | ESM2 residues, replacing ProtT5 | Full-reaction RXNFP | Convolution branches, bidirectional cross-attention, and regression head |

CatPred reads at most the first 2,047 residue vectors. MCKcat retains the first and last 500 vectors when a sequence exceeds 1,000 residues; these vectors are taken from the cached full-sequence representation. The other PLM baselines use all cached residues or their mean. UniKP follows its released 220-token SMILES rule, with a fixed extraction order and batch size of 100 to preserve the encoder’s batch-dependent positional encoding. For external TurNuP evaluation, CataPro and UniKP use the supplied substrate SMILES without removing components.

#### Training and learning-rate selection

We start from each released training configuration and apply the common source split and validation-RMSE selection rule. For every neural baseline, a seed-42 search compares *{*10^*™*4^, 3 *×* 10^*™*4^, 10^*™*3^, 3*×*10^*™*3^, 10^*™*2^ *}*; the selected learning rate is then fixed across repeated runs. Table 8 records the resulting settings. This protocol replaces PMAK’s released testbased checkpoint selection and CataPro’s cross-validation schedule with source-validation selection. PMAK retains its fixed training-record order. CatPred uses its mean– variance estimation loss and reports the predicted mean. DLKcat fits log_2_(*k*_cat_) after an exact change of base from the corrected labels, then converts predictions to log_10_ for evaluation. The remaining methods use the corrected log_10_ activities as their regression targets.

**Table 8.** Neural-baseline training settings. Learning rates are selected by source-validation RMSE from the common five-value grid. CatPred’s searched value is the peak learning rate. Each run retains its own best validation checkpoint.

| Method | Selected LR | Batch | Max. epochs | Loss | Schedule and retained settings |
|---|---|---|---|---|---|
| DLKcat | 10<sup>−3</sup> | 1 | 50 | MSE | Adam; weight decay 10<sup>−6</sup>; LR halved before epochs 10, 20, 30, 40, and 50 |
| CatPred | 10<sup>−3</sup> | 16 | 30 | MVE | Released training loop; initial and final LR equal 0.1 times the peak LR |
| CataPro | 10<sup>−4</sup> | 64 | 150 | RMSE | Adam; weight decay 0.01; dropout 0; early stopping with patience 20 and minimum improvement 0.001 |
| PMAK | 10<sup>−4</sup> | 1 | 300 | MSE | Adam; weight decay 10<sup>−5</sup>; fixed learning rate and full epoch budget |
| MCKcat | 10<sup>−4</sup> | 1 | 40 | MSE | Adam; weight decay 10<sup>−5</sup>; fixed learning rate and full epoch budget |

UniKP and TurNuP use the released tree-model hyperparameters and are fitted once on the combined 18,424 source training and validation records with seed 42. UniKP uses 100 ExtraTrees, unlimited tree depth, all input features at each split, and no bootstrap sampling, following the defaults of scikit-learn 1.7.2. TurNuP fits its enzyme and reaction regressors for 313 and 109 boosting rounds, respectively, with the released branch-specific XGBoost parameter blocks.

#### Repeated training and aggregation

The neural baselines use the same ten seeds as EnzRef (Table 9); the seed-42 search model is separate from these runs. Single-model scores report the arithmetic mean and population standard deviation (ddof = 0) of the ten run-level metrics. Ensemble scores are computed after averaging the ten predictions for each record in log_10_ space. UniKP’s result comes from its single fitted 100-tree regressor, while TurNuP averages its enzyme and reaction predictions with weights 0.5*/*0.5.

**Table 9.** Fixed training seeds. The same ordered seeds are used for the ten EnzRef runs and for each protein-only/EnzRef pair in the cross-PLM experiments.

| Run | Seed | Run | Seed |
|---|---|---|---|
| 1 | 1764580113 | 6 | 601323147 |
| 2 | 1479059094 | 7 | 1348470056 |
| 3 | 913210237 | 8 | 805792829 |
| 4 | 114158692 | 9 | 1685443149 |
| 5 | 723869210 | 10 | 885795228 |

#### Evaluation coverage

CatPred evaluates 2,048 of the 2,049 source held-out records because its D-MPNN cannot construct a graph for one reactant string, [H]. All other methods cover the full source held-out set. DLKcat follows the CatPred adapter’s vocabulary construction over the original CatPred training, validation, and test inputs. Its atom, bond, fingerprint, and sequence 3-mer vocabularies remain fixed during external evaluation, which excludes records containing unsupported symbols. DLKcat therefore covers 1,916 of the 2,746 external pair-unseen records and 696 of the 1,008 enzyme-unseen records. Its external scores use these supported subsets; the other methods cover both external subsets in full.

### C. Implementation Details

#### Chemical Semantic Decomposition (CSD)

Each molecular component is represented by a 2,048-bit, radius-3 Morgan fingerprint [42], computed with RDKit 2026.03.6. We use connectivity invariants, bond types, ring membership, chirality, and count simulation with thresholds *{*1, 2, 4, 8 *}*. The stored fingerprint is binary; count simulation encodes thresholded substructure counts into bits. Bitwise OR pools components within each reaction side, retaining reactant and product roles as separate chemical conditions. Both sides share a chemical encoder consisting of layer normalization, a 2,048 *→* 512 affine layer, GELU, dropout with probability 0.1, and a 512 *→* 256 affine layer.

#### RefCA and SASR

Let *D* denote the PLM feature dimension, with *D* = 1,280 for the default ESM2-650M backbone. RefCA has four query heads with 64-dimensional keys and queries and 256-dimensional values. The key map applies layer normalization without learned affine parameters and a bias-free *D →* 64 projection. The value map applies affine layer normalization, a *D →* 256 affine projection, and GELU. Keys and values are shared across roles and heads. Each head has its own bias-free 256 *→* 64 chemical-query projection and learned base query, shared between reactants and products. Base queries are initialized from *N* (0, 0.02^2^). Attention and the uniform reference use only valid residues.

For each head, SASR first computes the mean deviation and signed role difference of the two referenced responses, then applies separate layer-normalization, 256 *→* 16 affine, and GELU branches to these components. The branches have independent parameters across heads and components. Concatenating the eight outputs gives the 128-dimensional task representation. The predictor is a 128 *→* 128 affine layer, GELU, dropout with probability 0.1, and a scalar affine output. All layer normalizations except the key normalization have learned affine parameters. With ESM2-650M, the chemical encoder, RefCA, SASR, and predictor together contain 1,716,353 trainable parameters.

#### Optimization and model selection

We standardize log_10_(*k*_cat_) labels using the source-training mean and population standard deviation and optimize the mean squared error in Eq. 6. AdamW uses weight decay 10^*™*4^, batch size 512, and gradient clipping at an *ℓ*_2_ norm of 1. The maximum training length is 200 epochs, with five epochs of linear warmup followed by cosine learning-rate decay. A seed-42 search over *{*10^*™*4^, 3 *×* 10^*™*4^, 10^*™*3^, 3 *×* 10^*™*3^, 10^*™*2^*}* selects 10^*™*3^ by source-validation RMSE. This value is then fixed for the ten repeated training runs. Validation RMSE is measured after restoring predictions to the original log-activity scale. We retain a checkpoint when RMSE improves by more than 10^*™*4^ and stop after 20 epochs without such improvement.

#### Repeated training and compute

Each run starts from a separate random initialization with a seed from Table 9 and selects its own validation checkpoint. Default EnzRef training uses FP32 arithmetic and cached residue features on NVIDIA RTX 4090 GPUs. Four GPUs process independent runs, with one model per GPU at a time. The ten default runs stop after 69–129 epochs. Cross-PLM experiments use the same GPU type, with four or five independent runs scheduled concurrently.

### D. Cross-PLM Evaluation Protocol

#### Frozen feature extraction

Table 10 specifies the six frozen backbones. ESM2 features are extracted with fairesm 2.0.0 from the indicated final layer. For ProtT5-XL (1.2B encoder parameters), we use the encoder of <monospace>Rostlab/prot_t5_xl_uniref50</monospace> with Transformers 4.46.3 and its T5 tokenizer, taking the final hidden state. ProtT5 inputs use space-separated uppercase residues, with U, Z, O, and B replaced by X. Sequences longer than 1,022 residues are partitioned into nonoverlapping chunks of at most 1,022 residues. We encode chunks independently and concatenate their residue outputs in sequence order. Special tokens and padding are excluded. The protein-only input is the uniform mean over all retained residue vectors of the complete sequence, so longer sequences are represented by all their chunks.

**Table 10.** Frozen PLM features used in the paired comparison. *D* is the residue feature dimension. All features are inferred and stored in FP32 with TF32 disabled.

| Backbone | Output | *D* |
|---|---|---|
| ESM2-8M [7] | Layer 6 | 320 |
| ESM2-35M | Layer 12 | 480 |
| ESM2-150M | Layer 30 | 640 |
| ESM2-650M | Layer 33 | 1,280 |
| ProtT5-XL (1.2B) [8] | Final encoder layer | 1,024 |
| ESM2-3B | Layer 36 | 2,560 |

#### Protein-only predictor

The protein-only model maps the mean PLM vector through layer normalization, a *D →* 512 affine layer, GELU, dropout with probability 0.1, and a 512*→*256 affine layer. Its regression head receives this 256-dimensional vector concatenated with two 256-dimensional zero vectors in the chemical-input slots. The head has affine widths 768 *→* 512 *→* 128 *→* 1, with layer normalization, GELU, and dropout with probability 0.1 after each hidden affine layer. EnzRef uses CSD’s chemical conditions to query residue vectors through RefCA and recomposes the referenced responses with SASR. Across PLMs, the input dimension *D* changes while the remaining widths and the chemical inputs stay fixed.

#### Paired training and evaluation

Both predictors use identical source splits, corrected labels, and the ordered seeds in Table 9. Across all backbones, training follows the default ESM2-650M hyperparameters in Appendix C, including learning rate 10^*™*3^, batch size 512, and the same optimization and early-stopping schedules. For ESM2-650M, we reuse the default EnzRef models from Table 1; the protein-only predictor uses features extracted with the same encoder and extraction protocol. For the other backbones, frozen features are computed once and shared between the ten protein-only runs and ten EnzRef runs. Each run selects its checkpoint by source-validation RMSE. External evaluation applies the selected source-trained checkpoints directly, using the same feature-extraction rules without target-domain fitting.

### E. Conditional Activity-Difference Evaluation

#### Grouping and duplicate aggregation

The evaluations in Tables 2 and 4 use the same source held-out records and grouping rules. For shared-enzyme groups, we match complete amino-acid sequences exactly and distinguish reactions by the reaction identifiers in the frozen source dataset. We average the true and predicted log activities of duplicate records with the same enzyme and reaction identifier. An enzyme is eligible when at least two distinct reactions remain.

For shared-reaction groups, we canonicalize the reactant and product sides separately. Each side is split into molecular components, converted to canonical isomeric SMILES with RDKit, and sorted. We retain repeated components and preserve reaction direction, so exchanging reactants and products changes the reaction key. Within each canonical reaction, complete sequences identify enzymes; duplicate records for the same sequence are averaged in log space. A reaction is eligible when at least two distinct enzyme sequences remain.

#### Pair enumeration and aggregation

We enumerate all unordered pairs within each eligible group and apply Eq. 7. Both true and predicted differences use the same pair orientation; reversing that orientation leaves the squared error unchanged. The mean squared difference error is computed within each group, averaged equally over groups, and then square-rooted. Table 11 gives the resulting support. Groups containing only one distinct counterpart are excluded: 1,231 shared-enzyme groups and 1,583 shared-reaction groups. We compute the metric separately for each trained model and summarize the ten values using the mean and population standard deviation defined in Appendix B.

**Table 11.** Support for conditional activity-difference evaluation. Counts refer to eligible groups after duplicate aggregation. Each unordered pair contributes within its group.

| Shared input | Groups | Records | Pairs |
|---|---|---|---|
| Enzyme sequence | 317 | 818 | 864 |
| Canonical reaction | 190 | 466 | 458 |

### F. Held-Out Representation Analysis

#### Feature extraction

All comparisons use the same 2,049 held-out records. ESM-mean averages the frozen ESM2-650M residue vectors into a 1,280-dimensional sequence representation. These records contain 1,548 distinct enzyme sequences, so records sharing an enzyme have identical ESM-mean vectors. MCKcat’s 512-dimensional task representation concatenates the two interaction branches after residual connections, layer normalization, and proteinside max pooling. EnzRef’s 128-dimensional task representation concatenates the eight nonlinear SASR branch outputs. Both task representations are extracted immediately before the final prediction MLP from each of the ten trained models. Their run numbers follow Table 9.

#### Original-space cosine distributions

For each feature space, we normalize vectors to unit length and compute their dot products in float64. We include all 2,049 *×* 2,048*/*2 = 2,098,176 unordered record pairs, excluding self-pairs and retaining pairs that share an enzyme. Each task model yields its own cosine distribution. We use 200 equal-width bins on [*™*1, 1], normalize each histogram to unit area, and average the ten densities. The bands in Fig. 4b show the minimum and maximum density across models at each bin. The plotted range [*™*0.3, 1] retains every observed value. Table 12 summarizes the original-space statistics; ESM-mean has a single fixed representation and hence no between-run SD.

**Table 12.** EnzRef task features have lower average alignment. Mean cosine is summarized across runs; the range includes all pairs and runs. These are the feature spaces used in Fig. 4.

| Features | Dim. | Mean cosine | Observed range |
|---|---|---|---|
| ESM-mean | 1,280 | 0.900 | [0.374, 1.000] |
| MCKcat | 512 | 0.889 ± 0.006 | [0.487, 1.000] |
| EnzRef | 128 | 0.181 ± 0.027 | [−0.275, 0.999] |

**Table 13.**
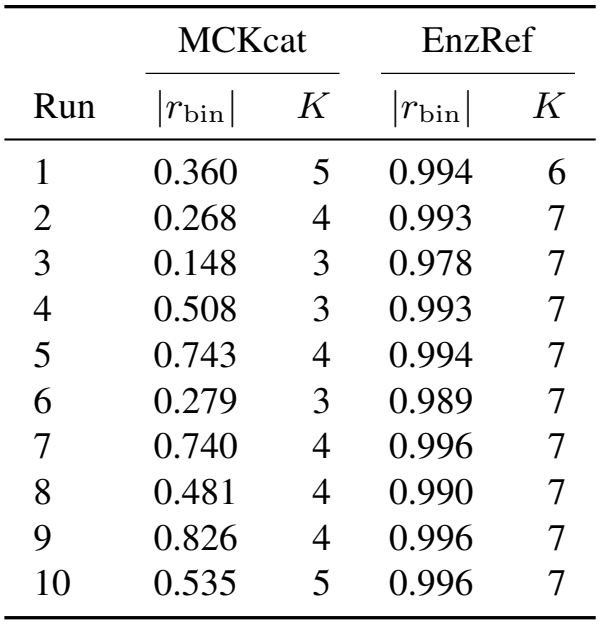
EnzRef shows more consistent activity progression across training runs. Absolute group-level correlation |*r*_bin_| and nondecreasing steps *K* out of seven, after orienting each projection. Run numbers match Fig. 8. All ten EnzRef guides and no MCKcat guides meet the full display criteria.

|  | MCKcat |  | EnzRef |  |
|---|---|---|---|---|
| Run | \|*r*<sub>bin</sub>\| | *K* | \|*r*<sub>bin</sub>\| | *K* |
| 1 | 0.360 | 5 | 0.994 | 6 |
| 2 | 0.268 | 4 | 0.993 | 7 |
| 3 | 0.148 | 3 | 0.978 | 7 |
| 4 | 0.508 | 3 | 0.993 | 7 |
| 5 | 0.743 | 4 | 0.994 | 7 |
| 6 | 0.279 | 3 | 0.989 | 7 |
| 7 | 0.740 | 4 | 0.996 | 7 |
| 8 | 0.481 | 4 | 0.990 | 7 |
| 9 | 0.826 | 4 | 0.996 | 7 |
| 10 | 0.535 | 5 | 0.996 | 7 |

#### Projection protocol

We fit each two-dimensional projection independently using its model’s precomputed cosine-distance matrix. The UMAP [50] views in Fig. 4 use 15 neighbors, minimum distance 0.1, spread 1, one worker, and random state 42. Task features use spectral initialization; ESM-mean uses random initialization because repeated sequence features cause the spectral initializer to fall back to a random start. The t-SNE [51] views in Fig. 9 use perplexity 30, early exaggeration 12, automatic learning rate, random initialization, 2,000 iterations, and random state 42, with the Barnes–Hut method and angle 0.5. Labels do not enter either projection fit. Each panel preserves its fitted coordinates and uses the same activity color scale; absolute axes and orientations are independent across panels. The label-based UMAP guides are described in Appendix G.

#### Sample-level activity association

Figure 4c shows the first three runs and panel (d) the first two in the shared training-seed order (Table 9). For each saved UMAP, we center its two-dimensional coordinates and obtain the first principal axis by singular value decomposition. Every sample is projected onto this axis. We orient its sign using the group-mean activity convention in Appendix G, then rescale the observed position range to [0, 1] separately for each projection. This affine rescaling preserves Pearson and Spearman correlations. Panel (d) plots all 2,049 positions against their true log-activities and reports their sample-level PCC (*r*) and SRCC (*ρ*). The line connects the mean positions and activities of eight equal-count groups along the oriented axis. These sample-level coefficients describe activity association within each projection; the guide statistics in Appendix G instead use eight group means. Figure 8 provides the complete ten-run comparison.

**Figure 8.**
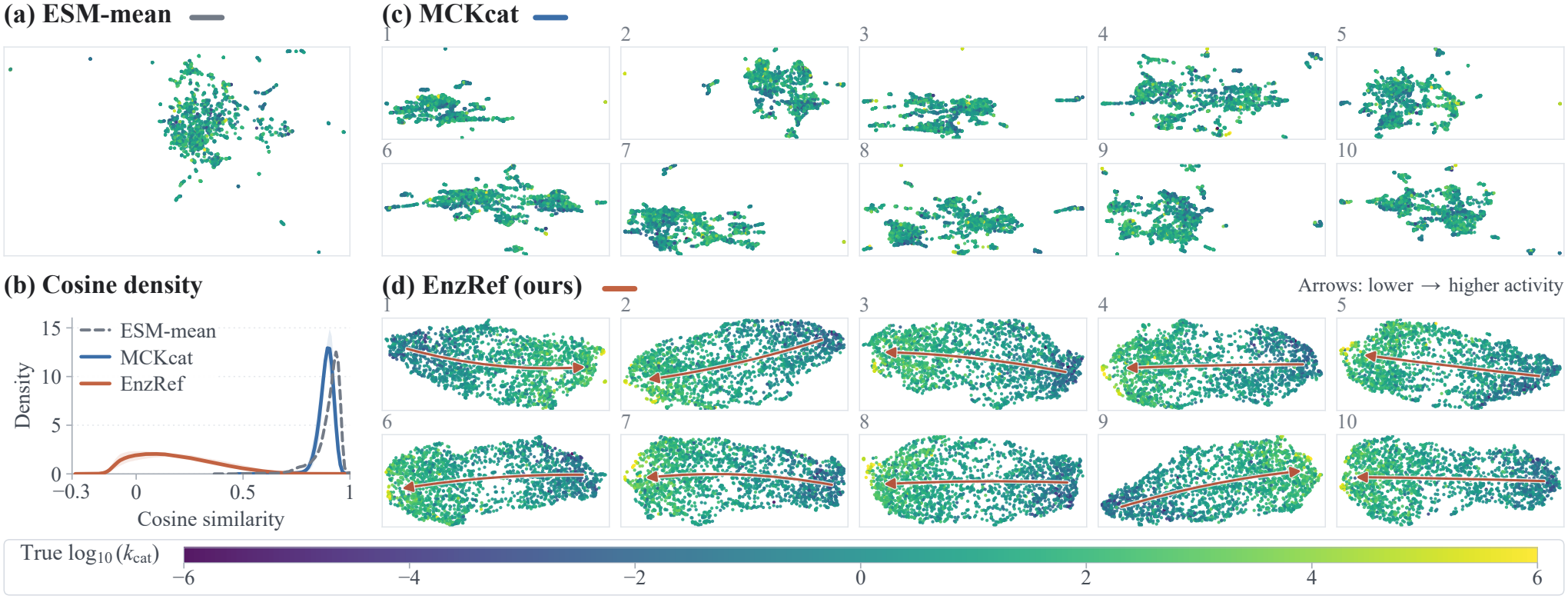
EnzRef’s projected activity progression persists across training runs. (a) Frozen ESM-mean UMAP. (b) Original-space cosine densities, with task-model means and min–max bands over ten runs. (c,d) All ten MCKcat and EnzRef UMAPs; matching numbers identify shared training seeds. Each panel uses the same 2,049 held-out records and activity color scale. Features alone determine the projections; activity labels orient and screen guides using the common rules in Appendix G.

**Figure 9.**
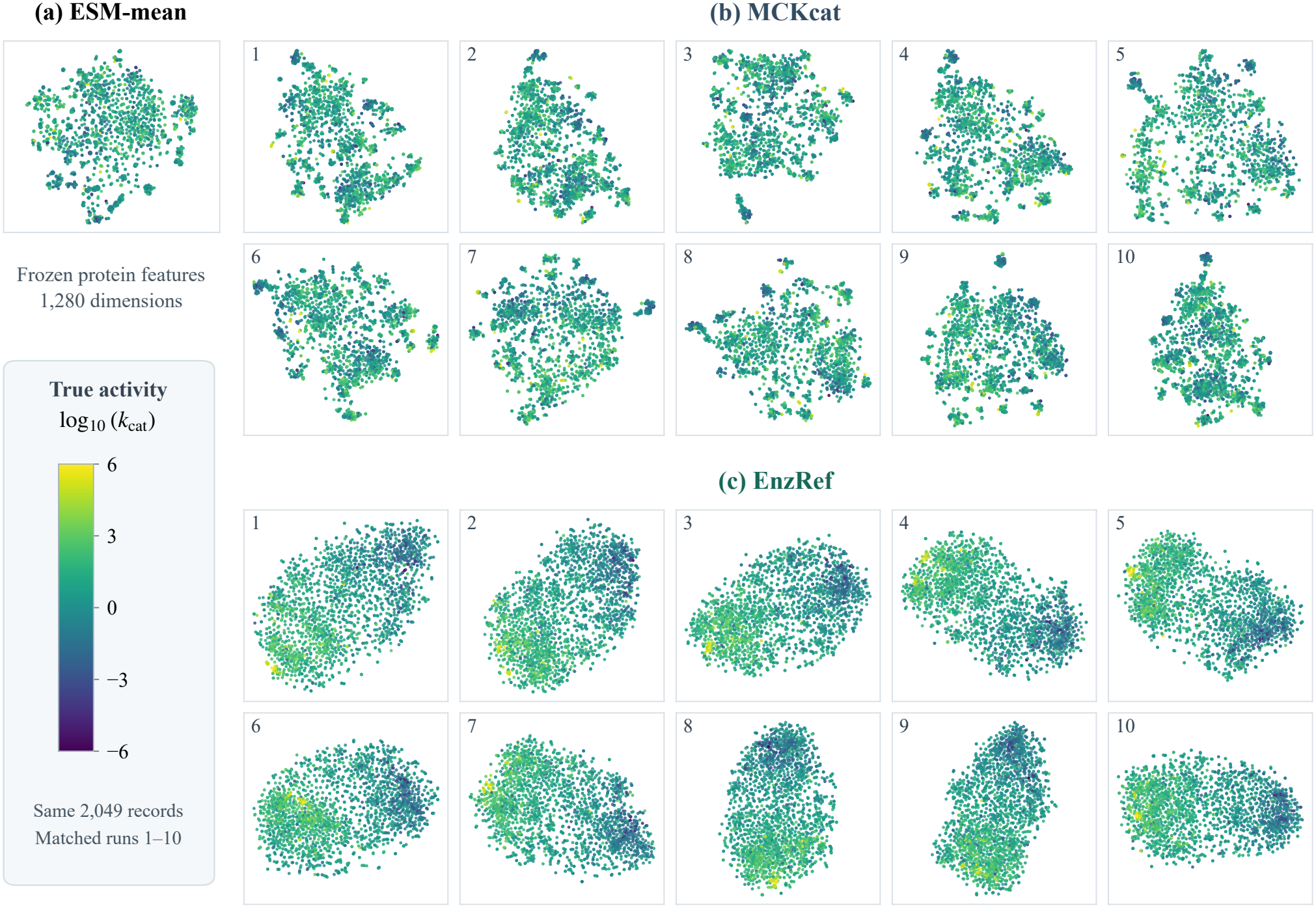
Activity gradients also appear in EnzRef’s t-SNE views. (a) Frozen ESM-mean. (b,c) All ten MCKcat and EnzRef task representations, with matching numbers identifying training seeds. Each panel contains the same 2,049 held-out records and uses the same true-activity color scale, marker size, and drawing order. Coordinates are fitted from features alone.

#### Complementary t-SNE views

EnzRef’s t-SNE projections show gradual activity changes within their main structures across all ten training runs. ESM-mean and MCKcat retain more separated local groups with greater color mixing. This qualitative pattern agrees with the UMAP observations under a different projection method. The original-space cosine analysis and the conditional-difference evaluation in Appendix E provide the accompanying geometric and predictive measurements.

### G. Activity Guides for UMAP Visualizations

We apply the same guide construction and display criteria to all ten MCKcat and ten EnzRef projections in Fig. 8. Each projection contains the same 2,049 held-out records and uses its own coordinate scale and orientation. The procedure uses the existing two-dimensional UMAP coordinates; the activity labels enter only the point colors, guide orientation, and display criteria. The PC1 positions in Fig. 4d use the same sign convention; their sample-level correlations are defined in Appendix F.

#### Grouping and direction

For each projection, we center its coordinates and obtain the two principal axes by singular value decomposition. We sort samples by the first principal coordinate and split them into eight contiguous groups of approximately equal size: one group of 257 samples and seven of 256. For group *b*, let (*s*_*b*_, *t*_*b*_) denote its centroid in the principal-axis basis and 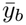 its mean true log_10_(*k*_cat_). We compute the Pearson correlation 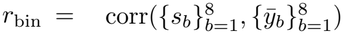 and orient the guide by its sign. If *r*_bin_ *<* 0, we reverse the group order for the following checks. Thus, |*r*_bin_| measures the association between group position and mean activity across the eight groups.

#### Geometric curve

We rescale the first principal coordinate so that the first and last group centers map to 1 and *™*1, respectively. A least-squares quadratic predicts the second principal coordinate from the first using the eight geometric centers. We map this curve back to the UMAP coordinates and draw it only between the first and last group-center positions. The curve is rendered as a quadratic Bézier path: for its endpoints *p*_0_, *p*_1_ and midpoint *p*_*m*_, the control point is 2*p*_*m*_ *™* (*p*_0_ + *p*_1_)*/*2. Labels determine its orientation and visibility, while its shape depends on coordinates alone.

#### Common display criteria

An arrow is shown when all four conditions hold: |*r*_bin_| *≥* 0.8; at least six of the seven adjacent group-mean differences are nonnegative in the oriented order; the last minus first group mean is at least 0.5 log units; and the curve remains close to observed samples. For the last condition, we evaluate 96 uniformly spaced first-coordinate positions along the curve. At each position, we calculate the nearest Euclidean distance to any observed point after dividing each coordinate difference by that co-ordinate’s full observed range. The maximum of these 96 distances must not exceed 0.08. These are descriptive display rules chosen during figure development and then applied unchanged to both models.

#### Per-run observations

All ten EnzRef runs meet the four display criteria, with |*r*_bin_| ranging from 0.978 to 0.996 and six or seven nondecreasing steps. MCKcat has three to five nondecreasing steps per run, below the six-step threshold. The median absolute group-level PCC is 0.993 for EnzRef versus 0.494 for MCKcat. These results show more consistent projected activity progression across EnzRef’s training runs.

### H. Additional Ablation Analyses

#### H.1. Full Factorial Ablation and Paired Interaction

Table 14 reports the complete 2 *×* 2 ablation of RefCA and SASR, including both conditional-difference tasks. Raw mean/diff. (b) is EnzRef w/o RefCA in Table 4, and Ref. sides (c) is EnzRef w/o SASR in Table 3. Raw sides (a) omits both reference centering and mean-and-contrast recomposition.

**Table 14.**
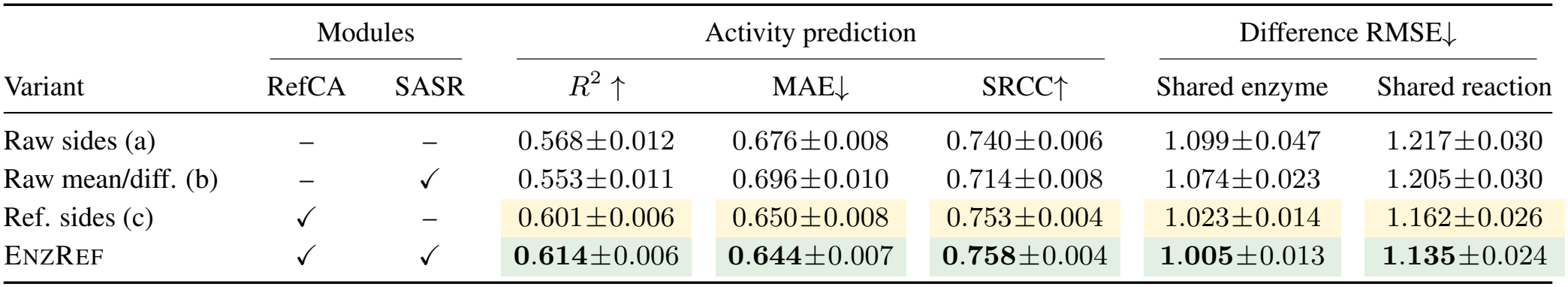
Full factorial ablation of RefCA and SASR. Source held-out entries are single-model means *±* population SD over ten matched runs. All variants retain CSD, frozen ESM2-650M features, and equal parameter counts. An unchecked RefCA uses uncentered cross-attention; an unchecked SASR encodes the two role readouts separately. Variants (b) and (c) are EnzRef w/o RefCA and EnzRef w/o SASR, respectively. Difference RMSE uses the groups in Table 2. **Green/bold** and yellow mark the best and second-best means.

|  | Modules |  | Activity prediction |  |  | Difference RMSE↓ |  |
|---|---|---|---|---|---|---|---|
| Variant | RefCA | SASR | *R*<sup>2</sup> ↑ | MAE↓ | SRCC↑ | Shared enzyme | Shared reaction |
| Raw sides (a) | – | – | 0.568±0.012 | 0.676±0.008 | 0.740±0.006 | 1.099±0.047 | 1.217±0.030 |
| Raw mean/diff. (b) | – | ✓ | 0.553±0.011 | 0.696±0.010 | 0.714±0.008 | 1.074±0.023 | 1.205±0.030 |
| Ref. sides (c) | ✓ | – | 0.601±0.006 | 0.650±0.008 | 0.753±0.004 | 1.023±0.014 | 1.162±0.026 |
| ENZREF | ✓ | ✓ | **0.614**±0.006 | **0.644**±0.007 | **0.758**±0.004 | **1.005**±0.013 | **1.135**±0.024 |

The overall prediction benefit of SASR’s mean-and-difference organization depends on enzyme referencing. With RefCA, moving from Ref. sides (c) to EnzRef raises mean single-model *R*^2^ from 0.601 to 0.614. Without RefCA, moving from Raw sides (a) to Raw mean/diff. (b) lowers it from 0.568 to 0.553, although both conditional-difference errors improve.

We quantify this dependence using the four variants’ paired prediction errors. Let *e*_*v,s*_ denote ordinary RMSE over all 2,049 source held-out records for variant *v* trained with seed *s*. The effect of mean-and-difference organization is 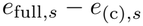 with RefCA and *e*_(b),*s*_*™ e*_(a),*s*_ without it. Their paired interaction is

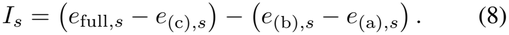

A negative interaction indicates that mean-and-difference organization is more favorable with enzyme referencing. We pair the ten training seeds across variants and compute a two-sided Student-*t* 95% confidence interval for the mean interaction, with nine degrees of freedom.

The mean interaction is *™*0.0316, with a 95% interval of [*™*0.0473, *™*0.0159]; nine of ten seeds yield a negative interaction. The interval summarizes variation across paired training seeds on the fixed held-out set.

#### H.2. Readout Geometry across All Variants

Figure 10 extends the main reference comparison to all four configurations in Table 14. Each configuration was trained independently. For each model, we concatenate the four heads of each chemical role into a 1,024-dimensional vector before nonlinear branch encoding. Variants with RefCA use the referenced readouts *r*_*s,h*_; variants without it use the raw readouts *u*_*s,h*_.

**Figure 10.**
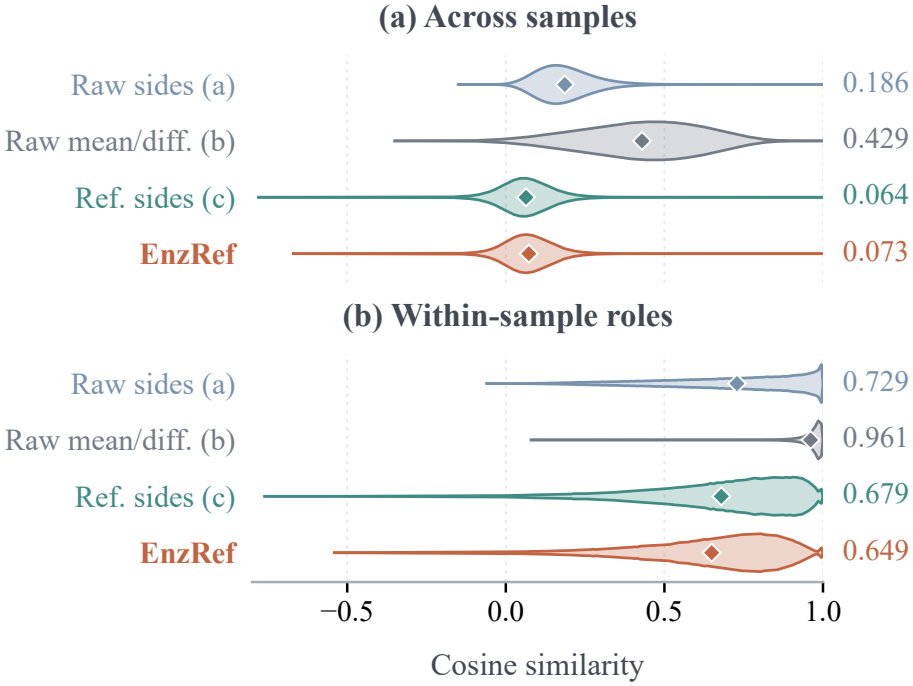
Readout geometry across the full RefCA/SASR ablation. Variants follow Table 14. Rows labeled (b) and (c) correspond to EnzRef w/o RefCA and w/o SASR, respectively. The panels show cross-sample and within-sample role cosine before nonlinear encoding. Violins show ten-run mean training-set densities with widths normalized separately; diamonds mark mean cosines.

For cross-sample comparison, we concatenate each record’s two role vectors into a 2,048-dimensional vector and evaluate all 137,555,991 unordered pairs among the 16,587 training records. Within-sample comparison uses one reactant–product cosine per record. Each distribution is computed separately for each of ten models, using identical sample order and training seeds across configurations.

Original counts use 4,000 equal-width bins on [*™*1, 1]. For display, we sum every 20 adjacent bins, normalize each model’s histogram to unit area, and average the ten densities. The violins mirror these histogram densities and normalize each violin to the same maximum width. Diamonds use means from the original moment statistics. Both panels retain the complete occupied support; only an empty tail is omitted.

Figure 11 displays the original density scale for the reference ablation, with straight segments between bin centers and constant extensions to the outer bin edges. Bands show the per-bin minimum and maximum densities across the ten training runs.

**Figure 11.**
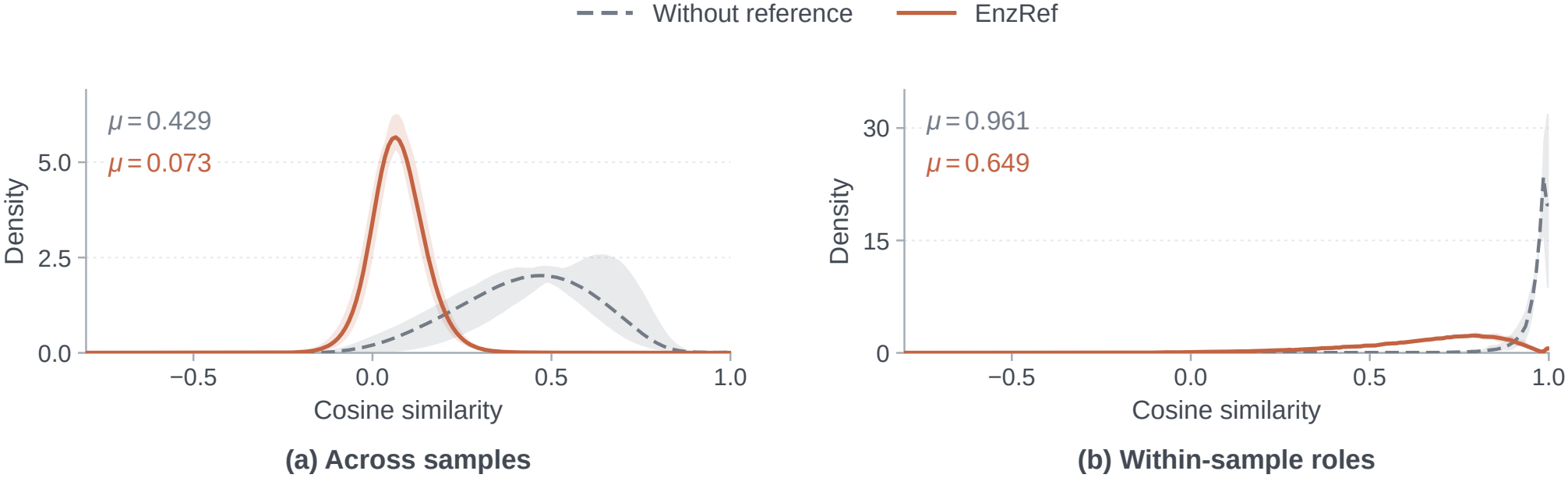
The reference changes readout alignment across training runs. EnzRef and the unreferenced variant (b) retain the same SASR architecture. (a) Cross-sample cosine of concatenated reactant/product readouts. (b) Within-sample reactant–product cosine. Curves and bands show ten-run mean densities and per-bin min–max ranges on the same 16,587 training records. Annotations give mean cosines.

### I. Efficiency Measurement

#### Parameter accounting

Figure 7 compares total single-model parameters after subtracting the common frozen ESM2-650M encoder. We count each shared parameter tensor once. The remaining count includes the task model and any other frozen neural encoders: RXNFP for PMAK and MCKcat, and MolT5 for CataPro. EnzRef and CatPred have no additional frozen neural feature extractor. Nonparametric fingerprints and molecular graph construction contribute no neural parameters.

The common ESM2 encoder contains 651,043,254 parameters. EnzRef adds 1,716,353 trainable parameters. CatPred adds 2,490,879 parameters, of which 304 are frozen. PMAK combines a 439,361-parameter task model with the 6,674,432-parameter RXNFP encoder, giving 7,113,793 parameters beyond ESM2. MCKcat uses the same RXNFP encoder with a 3,097,729-parameter task model, giving 9,772,161. CataPro combines a 572,161-parameter task model with a 109,628,544-parameter MolT5 encoder, giving 110,200,705. These counts define the bubble areas in Fig. 7; the frozen chemical encoders are counted even when their outputs are cached for timing.

#### Cached-input timing

We timed each of ten fixed check-points on the same ordered set of 2,048 records using FP32 on an RTX 5090 GPU, with AMP and TF32 disabled. Each checkpoint completed one full warmup pass followed by three measured passes. We took the median time within each checkpoint and report the mean and population standard deviation of these ten medians. Timing includes batch assembly from initialized caches, input transfer, the task-model forward pass, output retrieval, and implementation-specific postprocessing. Feature generation, model loading, cache initialization, and file writing are excluded. Times describe a single checkpoint, not the ten-model ensemble.

#### Batch selection and prediction consistency

We searched supported batch sizes for throughput using one fixed checkpoint, with prediction consistency as a selection requirement. EnzRef used batch 32. CataPro used batch 512. CatPred used batch 16, which exactly reproduced its saved evaluation predictions for all ten checkpoints; its faster batch-2 execution changed predictions and was excluded from the main comparison. The evaluated PMAK and MCKcat implementations supported only batch 1.

#### Accuracy and runtime sources

The plotted source *R*^2^ values and their standard deviations are taken from Table 1. They retain its evaluation coverage: 2,048 records for CatPred and 2,049 for the other models. Runtime was measured separately on the common 2,048 records. PMAK’s timed FP32 execution differs from its saved evaluation predictions by at most 0.005501 in log_10_(*k*_cat_). Its plotted *R*^2^ is retained from the saved predictions rather than recomputed on the timed execution path. Horizontal and vertical bars show the standard deviations of runtime and *R*^2^, respectively, across the ten runs.

### J. Complete Prediction Metrics

We report *R*^2^, RMSE, MAE, Pearson correlation (PCC), and Spearman rank correlation (SRCC) for the comparisons in Tables 1, 3, and 4 and Fig. 3. All metrics use targets and predictions in log_10_(*k*_cat_) space. PCC measures linear association, while SRCC measures agreement between predicted and observed activity ranks. We compute SRCC as Pearson correlation of the ranks, assigning average ranks to ties. Correlations are evaluated across all records in each evaluation set; the shared-enzyme and shared-reaction comparisons use the separate conditional-difference metric defined in Eq. 7.

#### Source prediction and zero-shot transfer

Tables 15–17 give the five metrics for the baseline comparison. EnzRef leads in prediction accuracy and activity rank agreement across all three evaluation sets in both evaluation modes. Repeated-run statistics and prediction aggregation follow Appendix B, with the coverage rules in Appendix B.

**Table 15.** Complete prediction metrics on the source held-out set (2,049 records). Single-model entries are mean *±* population SD over ten training runs. Prediction aggregation follows Table 1; **Green/bold** denotes the best value within each block. CatPred covers 2,048 records.

| Method | *R*<sup>2</sup> ↑ | RMSE↓ | MAE↓ | PCC↑ | SRCC↑ |
|---|---|---|---|---|---|
| *Independent training runs: mean ± SD* |  |  |  |  |  |
| DLKcat | 0.2460 ± 0.1268 | 1.2918 ± 0.1176 | 0.9675 ± 0.1113 | 0.4868 ± 0.1107 | 0.4658 ± 0.1136 |
| CatPred | 0.5439 ± 0.0096 | 1.0085 ± 0.0107 | 0.6953 ± 0.0105 | 0.7413 ± 0.0060 | 0.7321 ± 0.0084 |
| CataPro | 0.5074 ± 0.0069 | 1.0484 ± 0.0074 | 0.7658 ± 0.0088 | 0.7148 ± 0.0055 | 0.6885 ± 0.0071 |
| PMAK | 0.5187 ± 0.0122 | 1.0363 ± 0.0131 | 0.7461 ± 0.0105 | 0.7228 ± 0.0082 | 0.7046 ± 0.0098 |
| MCKcat | 0.5534 ± 0.0068 | 0.9982 ± 0.0077 | 0.7013 ± 0.0094 | 0.7469 ± 0.0045 | 0.7322 ± 0.0057 |
| ENZREF | **0.6138** ± 0.0057 | **0.9283** ± 0.0069 | **0.6445** ± 0.0070 | **0.7851** ± 0.0036 | **0.7579** ± 0.0045 |
| *Prediction aggregation* |  |  |  |  |  |
| DLKcat | 0.3149 | 1.2364 | 0.9317 | 0.5706 | 0.5611 |
| TurNuP | 0.4847 | 1.0724 | 0.7981 | 0.7185 | 0.7110 |
| UniKP | 0.5797 | 0.9685 | 0.6749 | 0.7623 | 0.7571 |
| CatPred | 0.5838 | 0.9634 | 0.6642 | 0.7645 | 0.7557 |
| CataPro | 0.5430 | 1.0099 | 0.7283 | 0.7370 | 0.7115 |
| PMAK | 0.5759 | 0.9728 | 0.7014 | 0.7633 | 0.7466 |
| MCKcat | 0.5989 | 0.9461 | 0.6590 | 0.7740 | 0.7617 |
| ENZREF | **0.6432** | **0.8923** | **0.6145** | **0.8022** | **0.7791** |

**Table 16.**
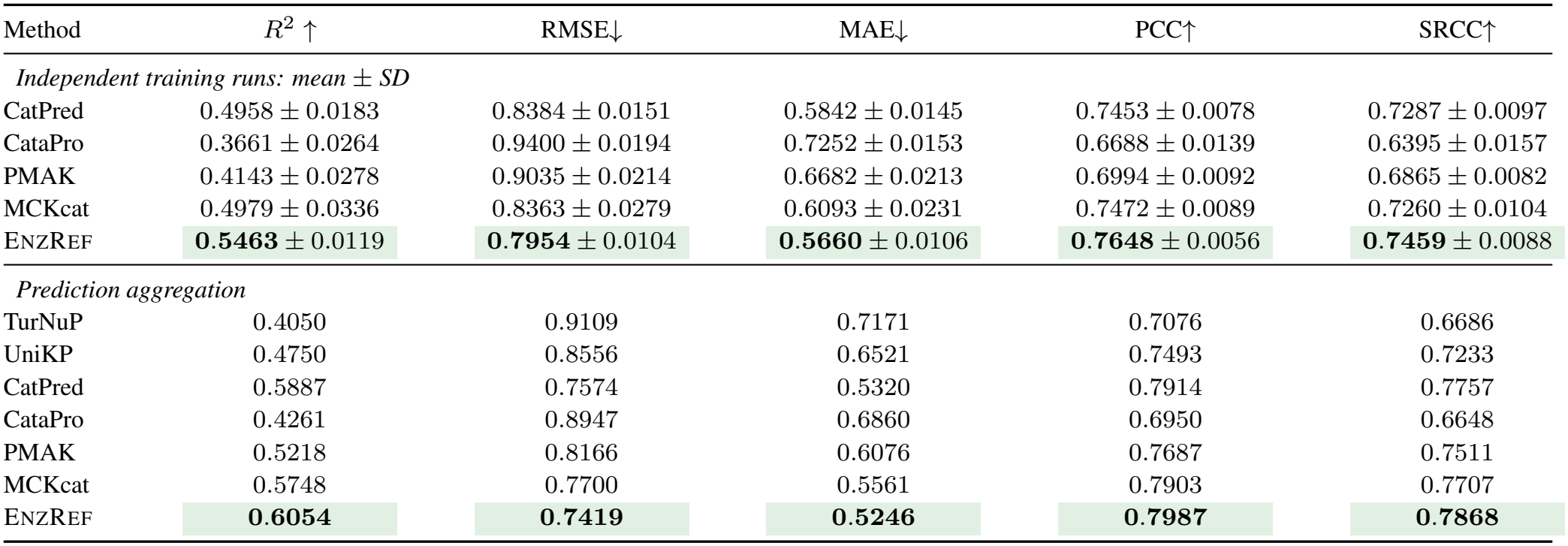
Complete prediction metrics on the external pair-unseen set (2,746 records). Single-model entries are mean *±* population SD over ten training runs. Prediction aggregation follows Table 1; **Green/bold** denotes the best value within each block. DLKcat is omitted because its frozen vocabulary does not cover the full subset (Appendix B).

**Table 17.** Complete prediction metrics on the external enzyme-unseen set (1,008 records). Single-model entries are mean *±* population SD over ten training runs. Prediction aggregation follows Table 1; **Green/bold** denotes the best value within each block. DLKcat is omitted because its frozen vocabulary does not cover the full subset (Appendix B).

| Method | *R*<sup>2</sup> ↑ | RMSE↓ | MAE↓ | PCC↑ | SRCC↑ |
|---|---|---|---|---|---|
| *Independent training runs: mean ± SD* |  |  |  |  |  |
| CatPred | 0.1720 ± 0.0431 | 1.0792 ± 0.0278 | 0.8225 ± 0.0203 | 0.5249 ± 0.0215 | 0.5032 ± 0.0268 |
| CataPro | 0.1655 ± 0.0299 | 1.0837 ± 0.0194 | 0.8341 ± 0.0174 | 0.5346 ± 0.0164 | 0.5052 ± 0.0196 |
| PMAK | 0.1474 ± 0.0508 | 1.0950 ± 0.0329 | 0.8551 ± 0.0291 | 0.5143 ± 0.0188 | 0.4829 ± 0.0226 |
| MCKcat | 0.2586 ± 0.0470 | 1.0210 ± 0.0323 | 0.7945 ± 0.0296 | 0.5869 ± 0.0214 | 0.5533 ± 0.0265 |
| ENZREF | **0.3099** ± 0.0156 | **0.9855** ± 0.0111 | **0.7557** ± 0.0114 | **0.6134** ± 0.0084 | **0.5840** ± 0.0132 |
| *Prediction aggregation* |  |  |  |  |  |
| TurNuP | 0.2348 | 1.0378 | 0.8328 | 0.5378 | 0.4833 |
| UniKP | 0.2477 | 1.0290 | 0.8052 | 0.5741 | 0.5280 |
| CatPred | 0.3548 | 0.9530 | 0.7326 | 0.6275 | 0.6102 |
| CataPro | 0.2264 | 1.0435 | 0.7993 | 0.5587 | 0.5251 |
| PMAK | 0.3008 | 0.9921 | 0.7819 | 0.6199 | 0.5814 |
| MCKcat | 0.3596 | 0.9494 | 0.7350 | 0.6535 | 0.6255 |
| ENZREF | **0.3945** | **0.9232** | **0.7045** | **0.6644** | **0.6520** |

#### Adaptation across PLMs

Tables 18–20 extend the paired comparison to all five metrics. EnzRef improves over each protein-only predictor across the three evaluation sets. The ESM2-650M EnzRef ensemble is reused from the main baseline comparison; the remaining ensembles follow the common training protocol in Appendix D.

**Table 18.**
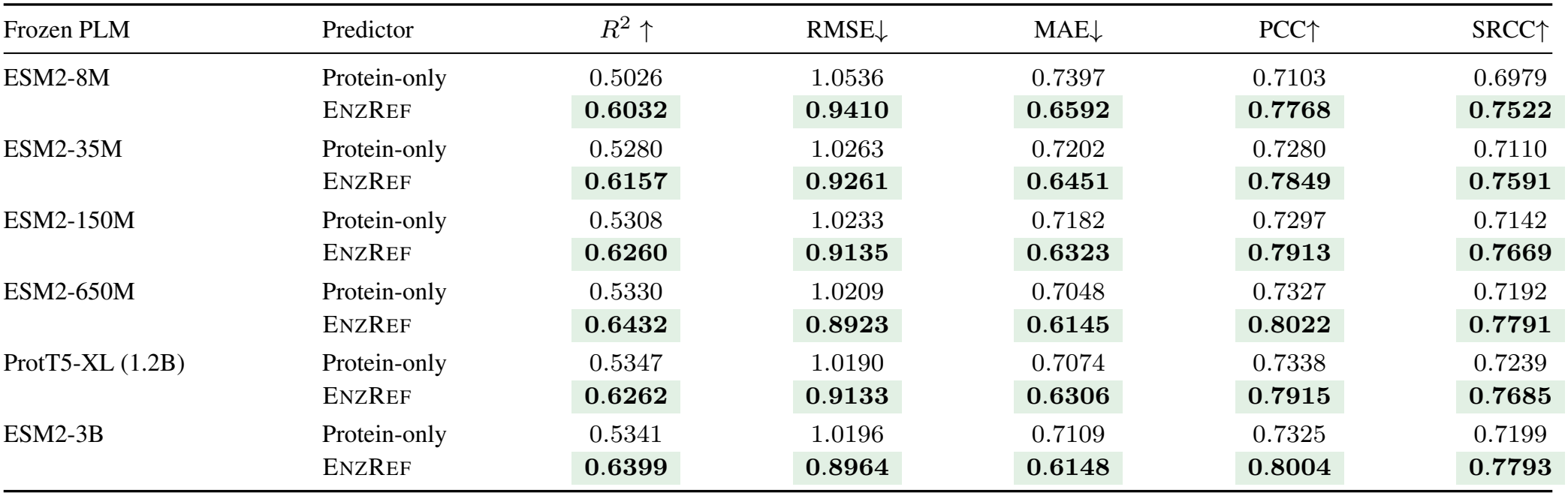
Complete cross-PLM metrics on the source held-out set. Scores use the ten-model prediction ensembles in Fig. 3. Each pair uses the same frozen PLM and training seeds; **Green/bold** marks the better value within each pair.

| Frozen PLM | Predictor | *R*<sup>2</sup> ↑ | RMSE↓ | MAE↓ | PCC↑ | SRCC↑ |
|---|---|---|---|---|---|---|
| ESM2-8M | Protein-only | 0.5026 | 1.0536 | 0.7397 | 0.7103 | 0.6979 |
|  | EnzRef | **0.6032** | **0.9410** | **0.6592** | **0.7768** | **0.7522** |
| ESM2-35M | Protein-only | 0.5280 | 1.0263 | 0.7202 | 0.7280 | 0.7110 |
|  | EnzRef | **0.6157** | **0.9261** | **0.6451** | **0.7849** | **0.7591** |
| ESM2-150M | Protein-only | 0.5308 | 1.0233 | 0.7182 | 0.7297 | 0.7142 |
|  | EnzRef | **0.6260** | **0.9135** | **0.6323** | **0.7913** | **0.7669** |
| ESM2-650M | Protein-only | 0.5330 | 1.0209 | 0.7048 | 0.7327 | 0.7192 |
|  | EnzRef | **0.6432** | **0.8923** | **0.6145** | **0.8022** | **0.7791** |
| ProtT5-XL (1.2B) | Protein-only | 0.5347 | 1.0190 | 0.7074 | 0.7338 | 0.7239 |
|  | EnzRef | **0.6262** | **0.9133** | **0.6306** | **0.7915** | **0.7685** |
| ESM2-3B | Protein-only | 0.5341 | 1.0196 | 0.7109 | 0.7325 | 0.7199 |
|  | EnzRef | **0.6399** | **0.8964** | **0.6148** | **0.8004** | **0.7793** |

**Table 19.**
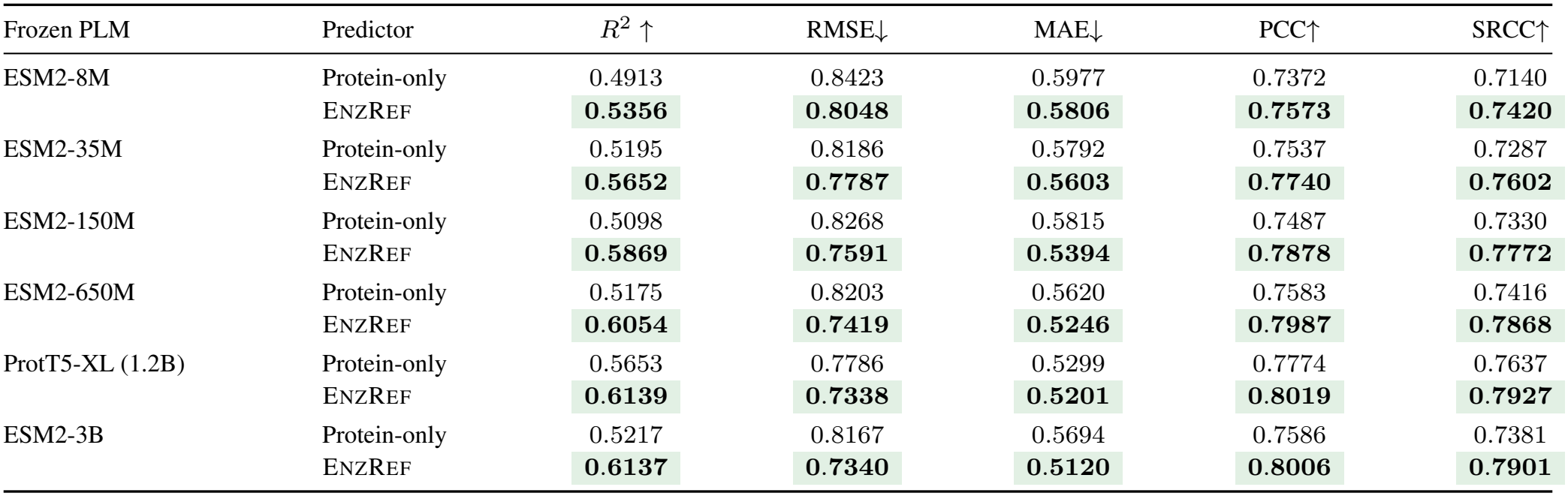
Complete cross-PLM metrics on the external pair-unseen set. Scores use the ten-model prediction ensembles in Fig. 3. Each pair uses the same frozen PLM and training seeds; **Green/bold** marks the better value within each pair.

| Frozen PLM | Predictor | *R*<sup>2</sup> ↑ | RMSE↓ | MAE↓ | PCC↑ | SRCC↑ |
|---|---|---|---|---|---|---|
| ESM2-8M | Protein-only | 0.4913 | 0.8423 | 0.5977 | 0.7372 | 0.7140 |
|  | EnzRef | **0.5356** | **0.8048** | **0.5806** | **0.7573** | **0.7420** |
| ESM2-35M | Protein-only | 0.5195 | 0.8186 | 0.5792 | 0.7537 | 0.7287 |
|  | EnzRef | **0.5652** | **0.7787** | **0.5603** | **0.7740** | **0.7602** |
| ESM2-150M | Protein-only | 0.5098 | 0.8268 | 0.5815 | 0.7487 | 0.7330 |
|  | EnzRef | **0.5869** | **0.7591** | **0.5394** | **0.7878** | **0.7772** |
| ESM2-650M | Protein-only | 0.5175 | 0.8203 | 0.5620 | 0.7583 | 0.7416 |
|  | EnzRef | **0.6054** | **0.7419** | **0.5246** | **0.7987** | **0.7868** |
| ProtT5-XL (1.2B) | Protein-only | 0.5653 | 0.7786 | 0.5299 | 0.7774 | 0.7637 |
|  | EnzRef | **0.6139** | **0.7338** | **0.5201** | **0.8019** | **0.7927** |
| ESM2-3B | Protein-only | 0.5217 | 0.8167 | 0.5694 | 0.7586 | 0.7381 |
|  | EnzRef | **0.6137** | **0.7340** | **0.5120** | **0.8006** | **0.7901** |

**Table 20.**
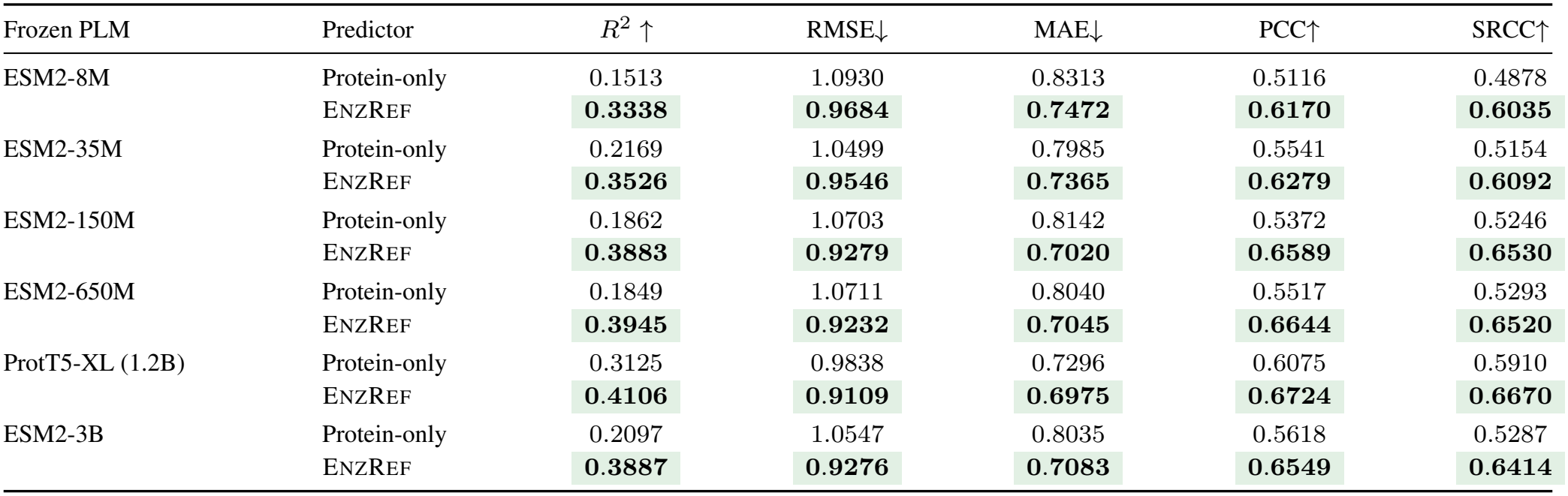
Complete cross-PLM metrics on the external enzyme-unseen set. Scores use the ten-model prediction ensembles in Fig. 3. Each pair uses the same frozen PLM and training seeds; **Green/bold** marks the better value within each pair.

| Frozen PLM | Predictor | *R*<sup>2</sup> ↑ | RMSE↓ | MAE↓ | PCC↑ | SRCC↑ |
|---|---|---|---|---|---|---|
| ESM2-8M | Protein-only | 0.1513 | 1.0930 | 0.8313 | 0.5116 | 0.4878 |
|  | EnzRef | **0.3338** | **0.9684** | **0.7472** | **0.6170** | **0.6035** |
| ESM2-35M | Protein-only | 0.2169 | 1.0499 | 0.7985 | 0.5541 | 0.5154 |
|  | EnzRef | **0.3526** | **0.9546** | **0.7365** | **0.6279** | **0.6092** |
| ESM2-150M | Protein-only | 0.1862 | 1.0703 | 0.8142 | 0.5372 | 0.5246 |
|  | EnzRef | **0.3883** | **0.9279** | **0.7020** | **0.6589** | **0.6530** |
| ESM2-650M | Protein-only | 0.1849 | 1.0711 | 0.8040 | 0.5517 | 0.5293 |
|  | EnzRef | **0.3945** | **0.9232** | **0.7045** | **0.6644** | **0.6520** |
| ProtT5-XL (1.2B) | Protein-only | 0.3125 | 0.9838 | 0.7296 | 0.6075 | 0.5910 |
|  | EnzRef | **0.4106** | **0.9109** | **0.6975** | **0.6724** | **0.6670** |
| ESM2-3B | Protein-only | 0.2097 | 1.0547 | 0.8035 | 0.5618 | 0.5287 |
|  | EnzRef | **0.3887** | **0.9276** | **0.7083** | **0.6549** | **0.6414** |

#### Chemical inputs and semantic recomposition

Table 21 extends the RXNFP and w/o CSD comparisons in Table 3 to all five metrics and includes Reactant-only, joint ECFP (OR), and product-zeroing controls. The w/o CSD variant is listed as Joint ECFP (Concat) to identify its chemical input. EnzRef achieves the best scores in both single-model and ensemble evaluation.

**Table 21.** Complete source held-out metrics for the chemical-input variants. The RXNFP and w/o CSD (Concat) comparisons in Table 3 are extended with Reactant-only, joint ECFP (OR), and product-zeroing controls. Single-model entries are mean *±* population SD over ten runs; ensemble scores use their averaged log-space predictions. **Green/bold** denotes the best value within each block.

| Variant | *R*<sup>2</sup> ↑ | RMSE↓ | MAE↓ | PCC↑ | SRCC↑ |
|---|---|---|---|---|---|
| *Independent training runs: mean ± SD* |  |  |  |  |  |
| Reactant-only ECFP | 0.5860 ± 0.0091 | 0.9611 ± 0.0106 | 0.6639 ± 0.0116 | 0.7696 ± 0.0054 | 0.7436 ± 0.0066 |
| Joint reaction (RXNFP) | 0.5532 ± 0.0078 | 0.9985 ± 0.0087 | 0.6990 ± 0.0061 | 0.7526 ± 0.0054 | 0.7265 ± 0.0065 |
| Joint ECFP (OR) | 0.5798 ± 0.0076 | 0.9684 ± 0.0088 | 0.6725 ± 0.0072 | 0.7671 ± 0.0046 | 0.7434 ± 0.0061 |
| Joint ECFP (Concat) | 0.5909 ± 0.0059 | 0.9554 ± 0.0069 | 0.6583 ± 0.0066 | 0.7731 ± 0.0030 | 0.7462 ± 0.0030 |
| Zeroed product ECFP | 0.5923 ± 0.0086 | 0.9538 ± 0.0101 | 0.6551 ± 0.0060 | 0.7726 ± 0.0051 | 0.7466 ± 0.0064 |
| ENZREF | **0.6138** ± 0.0057 | **0.9283** ± 0.0069 | **0.6445** ± 0.0070 | **0.7851** ± 0.0036 | **0.7579** ± 0.0045 |
| *Prediction ensemble* |  |  |  |  |  |
| Reactant-only ECFP | 0.6257 | 0.9139 | 0.6227 | 0.7917 | 0.7701 |
| Joint reaction (RXNFP) | 0.6006 | 0.9441 | 0.6521 | 0.7773 | 0.7546 |
| Joint ECFP (OR) | 0.6176 | 0.9238 | 0.6351 | 0.7875 | 0.7695 |
| Joint ECFP (Concat) | 0.6305 | 0.9080 | 0.6187 | 0.7948 | 0.7725 |
| Zeroed product ECFP | 0.6268 | 0.9126 | 0.6186 | 0.7922 | 0.7704 |
| ENZREF | **0.6432** | **0.8923** | **0.6145** | **0.8022** | **0.7791** |

The single-path ECFP variants use *f*_*R*_, *f*_*R*_ *∨ f*_*P*_, or [*f*_*R*_; *f*_*P*_] as the chemical encoder input for Reactant-only, Joint ECFP (OR), and Joint ECFP (Concat), respectively. Joint reaction (RXNFP) instead uses a joint reaction embedding [23]. All single-path variants retain enzyme-background referencing and encode one conditioned readout per attention head. Their four 16-dimensional branch outputs form a 64-dimensional prediction input. OR and Concat use chemical-encoder input widths of 2,048 and 4,096, respectively, with 1,685,441 and 2,738,113 trainable parameters; EnzRef has 1,716,353. Training follows the default protocol in Appendix C.

The additional Zeroed product ECFP control retains the full dual-side architecture but replaces product fingerprints with zeros throughout training and evaluation. This reduces mean *R*^2^ from 0.614 to 0.592, supporting the predictive value of sample-specific product chemistry.

Table 22 gives complete prediction metrics for all four RefCA/SASR configurations, including EnzRef w/o SASR from Table 3 and EnzRef w/o RefCA from Table 4. The corresponding conditional-difference results appear in Table 14. Mean-and-contrast encoding improves the referenced representations, whereas it reduces overall prediction performance for raw readouts.

**Table 22.**
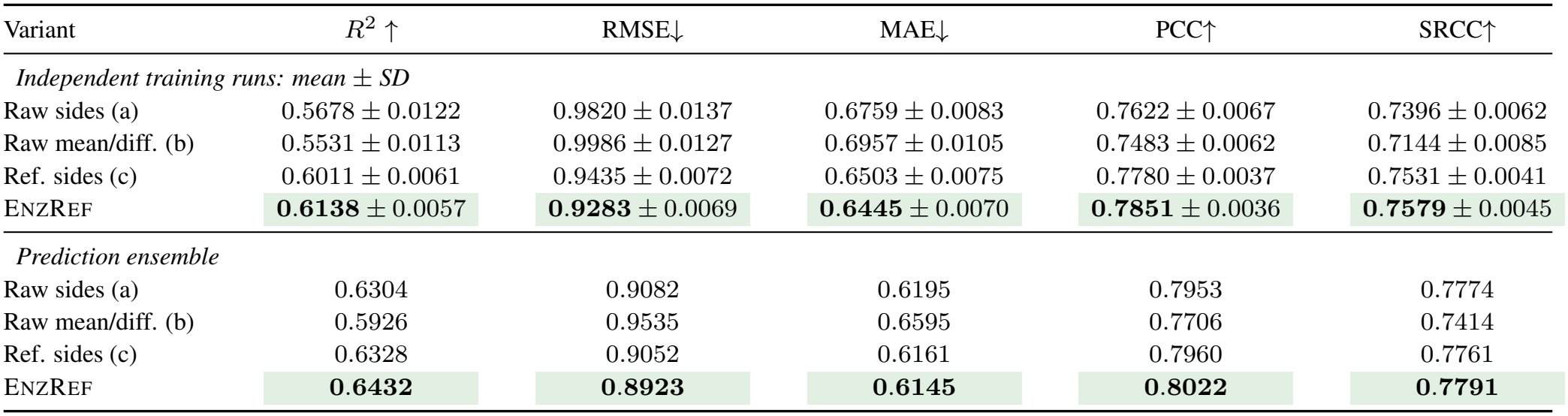
Complete source held-out prediction metrics for the RefCA and SASR variants. Rows follow the four configurations in Table 14; (b) is EnzRef w/o RefCA and (c) is EnzRef w/o SASR. Single-model entries are mean *±* population SD over ten runs; ensemble scores use their averaged log-space predictions. **Green/bold** denotes the best value within each block.

| Variant | *R*<sup>2</sup> ↑ | RMSE↓ | MAE↓ | PCC↑ | SRCC↑ |
|---|---|---|---|---|---|
| *Independent training runs: mean ± SD* |  |  |  |  |  |
| Raw sides (a) | 0.5678 ± 0.0122 | 0.9820 ± 0.0137 | 0.6759 ± 0.0083 | 0.7622 ± 0.0067 | 0.7396 ± 0.0062 |
| Raw mean/diff. (b) | 0.5531 ± 0.0113 | 0.9986 ± 0.0127 | 0.6957 ± 0.0105 | 0.7483 ± 0.0062 | 0.7144 ± 0.0085 |
| Ref. sides (c) | 0.6011 ± 0.0061 | 0.9435 ± 0.0072 | 0.6503 ± 0.0075 | 0.7780 ± 0.0037 | 0.7531 ± 0.0041 |
| ENZREF | **0.6138** ± 0.0057 | **0.9283** ± 0.0069 | **0.6445** ± 0.0070 | **0.7851** ± 0.0036 | **0.7579** ± 0.0045 |
| *Prediction ensemble* |  |  |  |  |  |
| Raw sides (a) | 0.6304 | 0.9082 | 0.6195 | 0.7953 | 0.7774 |
| Raw mean/diff. (b) | 0.5926 | 0.9535 | 0.6595 | 0.7706 | 0.7414 |
| Ref. sides (c) | 0.6328 | 0.9052 | 0.6161 | 0.7960 | 0.7761 |
| ENZREF | **0.6432** | **0.8923** | **0.6145** | **0.8022** | **0.7791** |

